# From implicit learning to explicit structural constraints for efficient protein language modeling

**DOI:** 10.64898/2026.03.17.712034

**Authors:** Lingdong Shen, Linlin Chao, Tang Liu, Qi Liu, Guowei Zhou, Hao Wang, Xinyue Dong, Tianhong Li, Xiaoming Zhang, Jinren Ni

## Abstract

Protein language models are typically pretrained on large sequence corpora, but unguided objectives can be inefficient at learning long-range structural dependencies. Here we introduce ProteinSage, an objective-level structure-constrained pretraining framework that uses AFDB-predicted residue-proximity priors during training while requiring only sequence inputs at inference. ProteinSage combines structure-guided masking of spatially proximal but sequence-distant residues with structural causal learning that predicts directional residue-pair dependencies. Across structure-aware and general protein benchmarks, ProteinSage achieves competitive or improved performance with substantially less training data and computation. We further applied it to metagenomic-scale screening of multi-pass transmembrane proteins and prioritized six rhodopsin-like candidates lacking prior characterization, which were validated as light-driven proton pumps. Together, our work shows that encoding predicted structural priors directly into pretraining objectives can improve the efficiency and transferability of protein language models without requiring structural inputs at inference.

## 1 Introduction

Recent advances in protein language models (PLMs) have substantially improved our ability to extract evolutionary, structural and functional information directly from amino acid sequences. Large-scale PLMs have enabled broad applications in structure prediction, variant effect estimation, functional annotation and protein design. Most existing PLMs are trained with token-level sequence objectives, including masked-language modeling (MLM) with standard random masking (SRM), as used in ESM-2 [1], ESM-3 [2] and xTrimoPGLM [3], or autoregressive next-token prediction, as used in ProGen [4] and ProtGPT2 [5]. Although scalable and effective, these objectives do not explicitly specify which residue dependencies are structurally informative, so long-range contacts and fold-consistent interactions must emerge implicitly from sequence statistics during pretraining [6].

Such implicit learning can be inefficient. When pretraining objectives do not prioritize structure-bearing positions, learning signals are broadly distributed across sequence sites, requiring larger protein corpora [7, 8] and greater compute to recover long-range dependencies. This unguided allocation of learning capacity may reduce the efficiency with which structural regularities are internalized, especially in redundant or weakly annotated data [9]. At scale, these costs also have environmental implications (Fig. 1a), as billion-parameter PLMs trained on multi-trillion-token corpora incur non-trivial carbon and water footprints [10]. Structural bioinformatics suggests a more targeted alternative. Structural and functional constraints are unevenly distributed along protein sequences (Fig. 1b). Co-evolving residues are often spatially proximal and associated with structural contacts, as shown by mutual-information analyses [11], direct-coupling methods [12], and mutational-coupling studies [13]. Proteome-scale analyses further show that structure–evolution couplings are enriched in active sites, ligand-binding pockets and interfaces [14], overlap residue contact maps [15], and form conserved functional sectors [16]. These observations suggest that a small subset of spatially proximal and evolutionarily constrained residue pairs carries disproportionate structural and functional signal, motivating pretraining objectives that emphasize these positions rather than relying only on uniformly sampled sequence-token prediction.

**Fig. 1.**
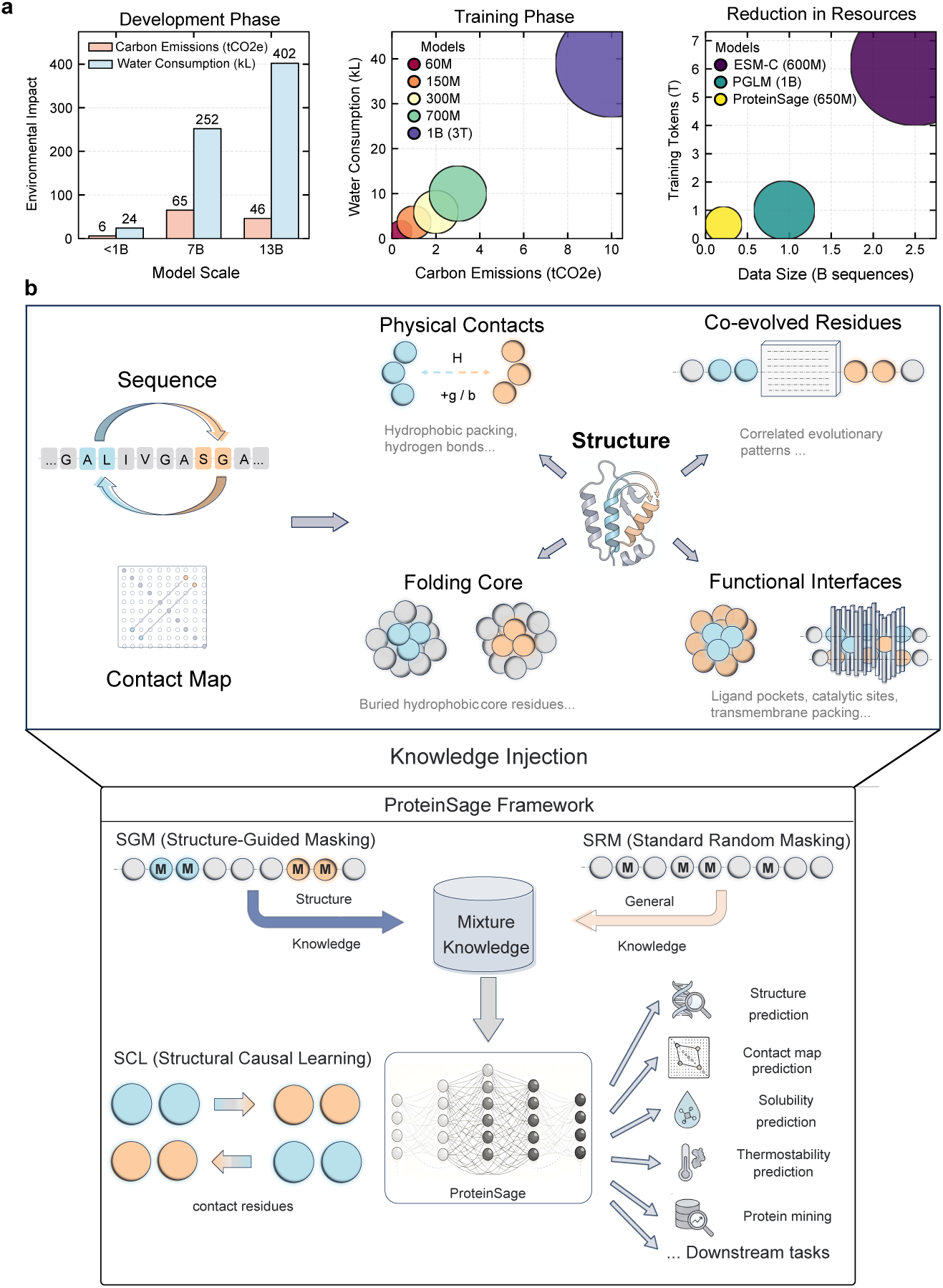
Training efficiency and overview of the ProteinSage framework. **(a), Environmental, computational and data efficiency of model training.** Estimated environmental costs are summarized across model scales. During development, representative (*<* 1)B, 7B and 13B models require 6/24, 65/252 and 46/402 tCO_2_e/kL, respectively. During training, the estimated cost increases from 0.4/1.6 tCO_2_e/kL for 60M models to 10.0/39.0 tCO_2_e/kL for 1B models. In terms of training resources, ProteinSage 650M uses 0.214B sequences and 0.5T tokens, compared with approximately 2.5 B/6.2T for ESM-C 600M and 0.94B/1.0T for PGLM 1B. Circle areas in the scatter plots indicate the relative combined scale of the two plotted quantities. **(b), Structure-constrained pretraining in ProteinSage.** ProteinSage uses predicted structural constraints to guide pretraining. SGM preferentially masks structurally important residues; SRM preserves standard random masking, and SCL models spatially proximate residue pairs to capture long-range dependencies. The resulting sequence-only pretrained model transfers to diverse downstream tasks.

The advent of AlphaFold and AFDB [17, 18] has made predicted structures available at proteome scale, enabling several structure-aware PLMs. Existing approaches extract contacts from PLM attention maps [19], distill fold-level information into sequence encoders [20], inject structural modules or adapters into pretrained transformers [21], use Foldseek-derived structural alphabets or structure-aware vocabularies [22, 23], quantize structures into discrete tokens [24], or align sequence, structure and functional text representations through contrastive learning [25]. These studies demonstrate the value of explicit structural information, but they generally introduce structure as an additional input, module, token vocabulary or aligned modality. At AFDB or metagenomic screening scale, such structure-dependent strategies can impose substantial computational, storage and input-processing costs because structural representations must be generated, retrieved or encoded for large numbers of candidate proteins (Extended Data Fig. 1a). This raises a central question: can predicted structural priors be transferred into the pretraining objective itself, so that structural information shapes representation learning during training while inference remains sequence-only?

Here we introduce ProteinSage, an objective-level structure-constrained PLM framework that uses predicted structural priors during pretraining but requires only amino acid sequences during inference. ProteinSage constructs residue-neighborhood graphs from AFDB-predicted structures and converts spatially proximal, sequence-distant residue pairs into pretraining signals. Structure-guided masking (SGM) preferentially masks structurally informative residues; standard random masking (SRM) preserves general sequence modeling capacity; and structural causal learning (SCL) introduces directed paired-residue prediction to capture long-range, fold-consistent dependencies. By moving structural information from the input space into the training objective, ProteinSage improves structural representation learning without requiring per-protein structure prediction or structural tokenization during downstream use.

Under matched pretraining protocols, ProteinSage achieves improved structural reasoning relative to a similarly sized ESM-C model while using more than an order of magnitude fewer training entries and tokens (Fig. 1a). Across structure-informed and general protein modeling benchmarks, ProteinSage shows competitive or improved performance with substantially lower training-resource requirements. We further develop ProteinSage-Miner for large-scale metagenomic screening of microbial rhodopsin candidates, whose seven-transmembrane architectures impose strong structural and functional constraints [26, 27]. It provides a complementary, structure-aware ranking signal for prioritizing low-sequence-similarity candidates for experimental validation. Together, these results establish ProteinSage as a blueprint for objective-level structure-constrained pretraining, demonstrating that predicted structural priors can be distilled into pretraining objectives to improve the data efficiency and transferability of PLMs without requiring structural inputs at inference.

## 2 Results

### 2.1 ProteinSage learns sequence-only representations through structure-constrained pretraining

*ProteinSage* is an objective-level structure-constrained pretraining framework that uses predicted residue-proximity priors during training while retaining sequence-only inference. It combines structure-guided masking (SGM), which preferentially masks spatially proximal but sequence-distant residue pairs, with structural causal learning (SCL), which converts these paired residues into directional dependency-prediction targets. Together, these objectives move structural information from an auxiliary input or post-hoc signal into the pretraining objective itself.

ProteinSage uses a Transformer encoder but changes how pretraining signals are allocated. SGM biases masking toward residue pairs linked by structure-derived spatial constraints (Methods 4.1.2; Fig. 1b), whereas SCL requires the model to predict one residue from its paired structural neighbor and surrounding sequence context (Methods 4.1.3). This design turns long-range residue dependencies into explicit training targets, rather than leaving them to be recovered only indirectly from token reconstruction.

By concentrating supervision on predicted residue-proximity relationships, ProteinSage improves the efficiency of structure-aware representation learning under reduced training budgets. Because predicted structures are used only to construct pretraining objectives, ProteinSage does not require structural inputs during downstream inference. The resulting sequence representations transfer across structure-informed and general protein modeling benchmarks, achieving competitive or improved performance without increasing model size. To evaluate downstream utility, we further develop **ProteinSage–Miner** (Methods 4.2), a lightweight adapter-based framework for prioritizing protein families with characteristic architectures, such as seven-transmembrane microbial rhodopsin-like proteins. ProteinSage–Miner combines frozen ProteinSage embeddings with a six-layer Transformer adapter and a classification head, enabling efficient metagenomic-scale candidate prioritization with a small number of trainable parameters.

### 2.2 ProteinSage improves structural reasoning

We next evaluated whether ProteinSage improves structural reasoning. We evaluated this property on four structural benchmarks: CAMEO [28], CASP15 [29], CASP14 [30], and Recent [1]. ProteinSage was compared with two strong baselines, PSL [9] and ESM-C [31], both pretrained on substantially larger datasets and longer training schedules. To isolate representation quality, all encoders were frozen and only a shallow structural prediction head was trained on top of the embeddings.

As shown in Fig. 2b(i,ii), ProteinSage reaches convergence using approximately 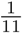 of ESM-C’s clustered training entries and 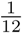 of its training tokens, while achieving stronger structural reasoning performance. Final-layer attention visualizations (Fig. 2a) show that ProteinSage places more attention on long-range, fold-consistent residue interactions, whereas ESM-C shows more diffuse attention and misses several reference structural-contact patterns. These observations suggest that structure-constrained pretraining helps redirect model capacity toward structure-relevant long-range dependencies.

**Fig. 2.**
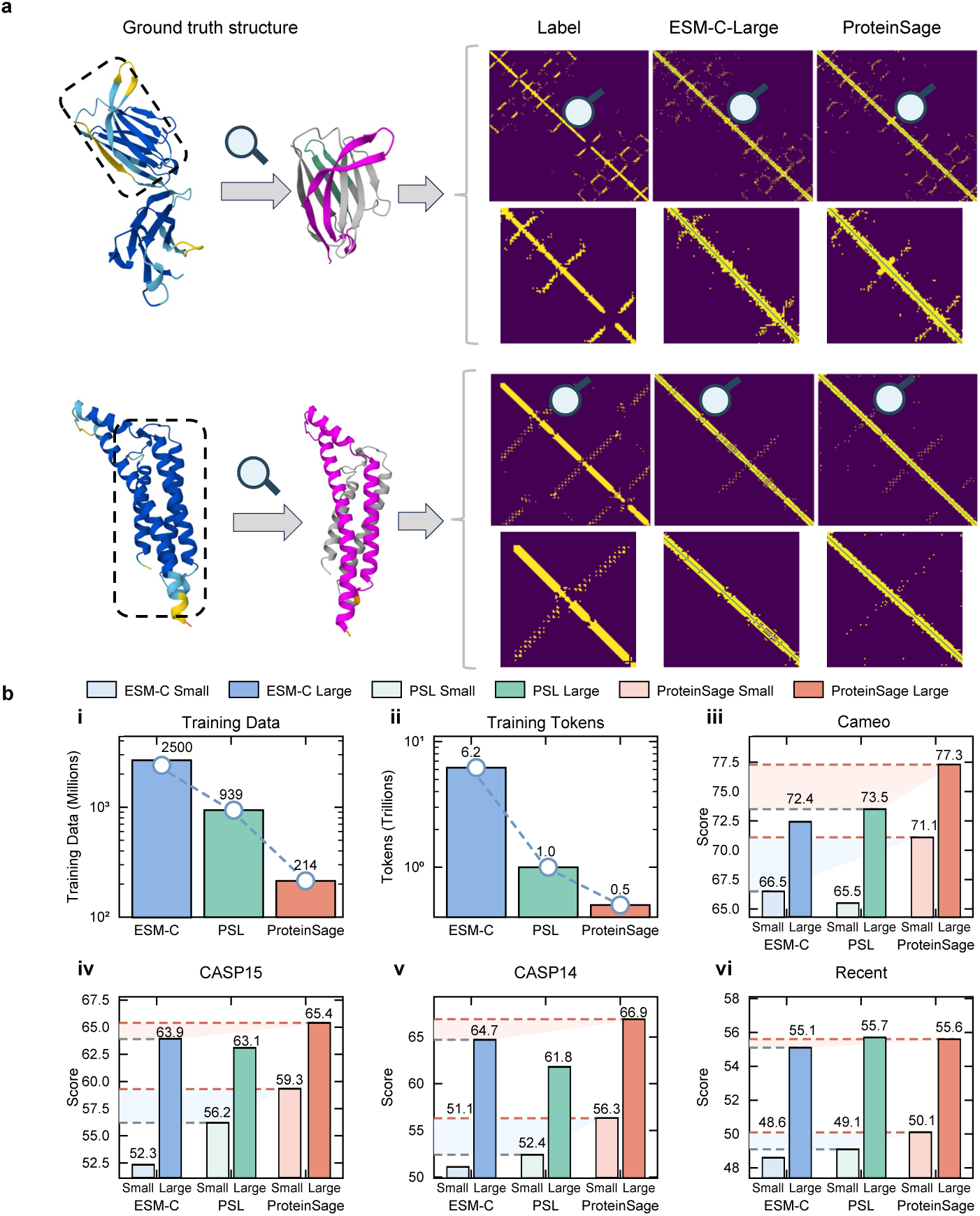
Unsupervised structural learning with ProteinSage. **(a)**, Heatmaps of final-layer pairwise attention (brighter indicates stronger coupling) for ProteinSage and ESM-C. ProteinSage recovers fold-consistent long-range interaction patterns indicative of contacts, whereas ESM-C misses several reference structural-contact patterns. **(b)**, Unsupervised structural prediction performance. (i,ii) Pretraining-entry counts and total training-token counts for three comparator model families, showing that ProteinSage achieves strong performance with fewer training entries and tokens. (iii–vi) Performance on four contact-map prediction benchmarks. Shaded regions indicate model-size–dependent performance gaps (blue, small models; red, large models); ProteinSage ranks first on three of the four benchmarks.

Across the four benchmarks (Fig. 2b(iii–vi)), ProteinSage achieves top or near-top performance across both model-size regimes. On CAMEO, ProteinSage ranks first for both small and large models, exceeding the second-best method by 4.6 and 3.8 points, respectively. Similar trends are observed on CASP15 and CASP14, where ProteinSage leads by 3.1–3.9 points in the small-model setting and by 1.5–2.2 points among large models. On Recent, ProteinSage maintains an advantage over ESM-C in the small-model regime and remains competitive with PSL despite a substantially smaller training footprint. To further examine potential overlap with AlphaFold-derived pretraining resources, we quantified sequence and structural similarity between the structural-reasoning benchmarks and AFDB v4 (Extended Data Fig. 2). This analysis identified conservative AFDB-distant subsets, including 149 proteins with low sequence similarity, 115 with low structural similarity and 56 with both low sequence and structural similarity. On these low-similarity subsets (Extended Data Fig. 3), ProteinSage also achieved the highest contact precision across sequence-distant, structure-distant and combined sequence–structure-distant proteins, with a smaller performance drop than ESM-C 600M and ESM-2 650M under the most stringent setting.

Together, these results indicate that objective-level structural priors improve the efficiency with which PLMs acquire transferable structural-reasoning capabilities, rather than relying primarily on larger data scale or longer training.

### 2.3 Structure-aware representations improve downstream performance

The transferability of ProteinSage representations was further evaluated across supervised protein modeling tasks. We compared ProteinSage with representative protein language models spanning encoder-style masked language models (ESM-2 [1], ESM-C [31], PGLM [3]), an encoder–decoder MLM (ProtT5 [32]), an autoregressive model (ProtGPT2 [5]), and a diffusion-based model (DPLM [33]) across eight supervised downstream tasks (Fig. 3a–h). These baselines cover major pretraining paradigms with public checkpoints. The task suite includes structural perception (fold classification [34], secondary-structure prediction [35], contact-map prediction [36]), functional and interaction phenotypes (antibiotic resistance [37], human protein–protein interaction [38], subcellular localization [39]), and biophysical properties (thermostability [40], solubility [41]). All models were evaluated under a unified supervised fine-tuning protocol to reduce implementation-specific confounders.

**Fig. 3.**
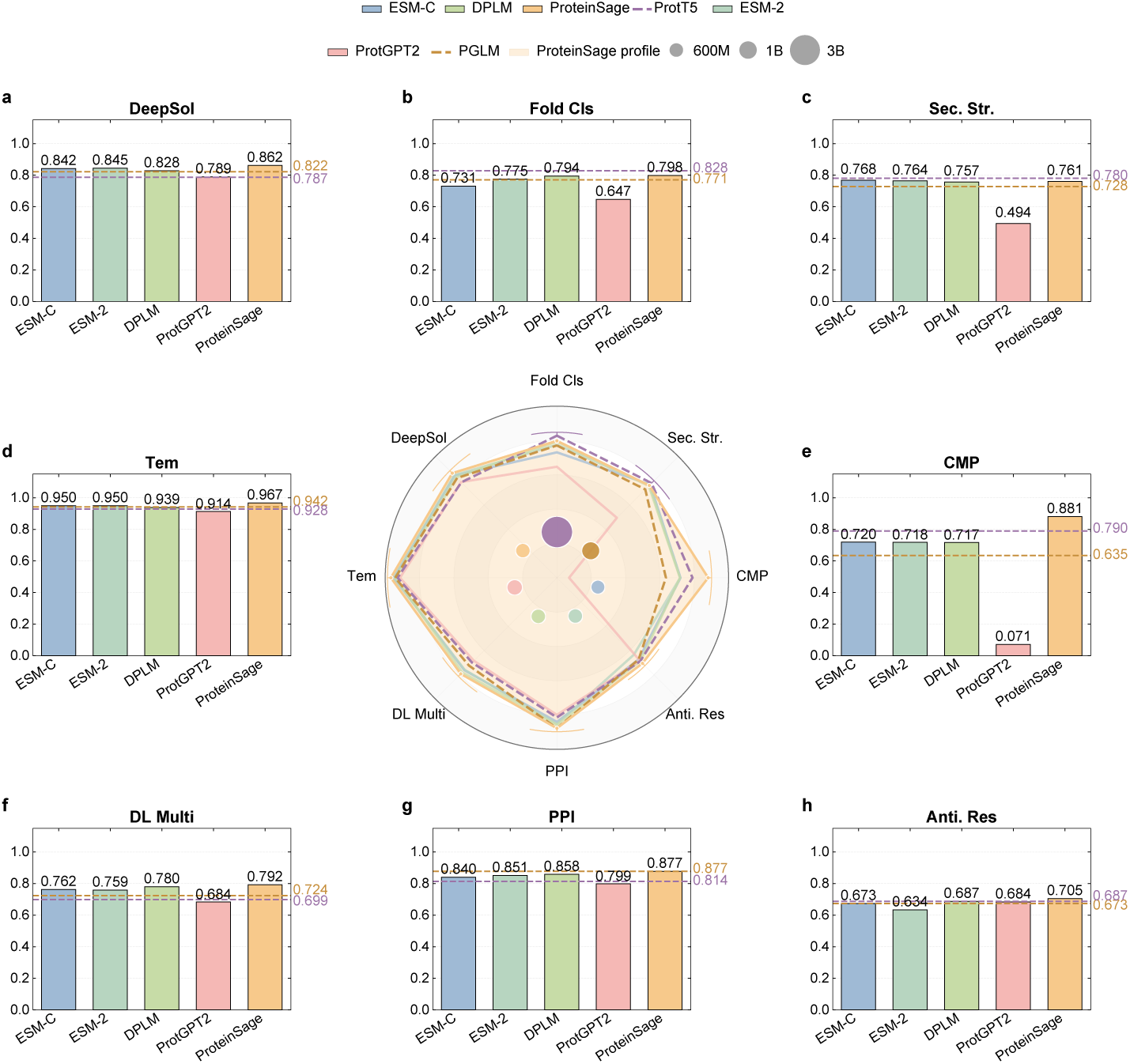
Supervised fine-tuning performance across protein tasks. Model parameter scales are illustrated in the central radar plot, where each scatter point represents a protein language model and marker area is proportional to its parameter count. The shaded polygon denotes the ProteinSage performance profile. The comparison covers encoder-style masked language models (ESM-2, ESM-C, PGLM), an encoder–decoder model (ProtT5), an autoregressive model (ProtGPT2), and a diffusion-based model (DPLM). Surrounding bar charts show supervised fine-tuning performance across eight downstream tasks: (a) solubility prediction, (b) fold classification, (c) secondary-structure prediction, (d) thermostability prediction, (e) contact map prediction, (f) subcellular localization, (g) human protein–protein interaction (PPI) prediction, and (h) antibiotic-resistance classification. Across these benchmarks, ProteinSage achieves the strongest overall performance, outperforming models of comparable scale (*<*1B parameters) on most tasks and matching or exceeding substantially larger models on several structure-related tasks, indicating efficient parameter utilization and strong downstream transfer under supervised fine-tuning.

With 650 M parameters, ProteinSage attains the highest mean performance across tasks (0.830), despite being comparable in size to ESM-2 and DPLM and smaller than ProtT5 (3 B) and PGLM (1 B). It exceeds DPLM (0.795) by +0.035 and ProtT5 (0.789) by +0.041, with larger margins over ESM-2 (+0.043), ESM-C (+0.045), and PGLM (+0.059). At the task level, ProteinSage leads or ties on six of eight benchmarks, with gains on contact-map prediction, antibiotic resistance, human PPI prediction, subcellular localization, thermostability, and solubility. Among similarly sized models (*<*1 B parameters), ProteinSage performs best on ^7^ tasks and matches or exceeds larger baselines on 6 of 8 tasks, while using approximately 4.6× fewer parameters than ProtT5.

These results suggest that objective-level structural priors improve not only contact-oriented reasoning but also broader sequence–structure–function transfer. The strongest gains occur on tasks where long-range residue dependencies, interaction interfaces, or stability-related constraints are expected to be informative, including contact prediction, PPI, thermostability and solubility. Rather than showing uniform dominance across all tasks, ProteinSage provides a parameter-efficient representation that is particularly effective for structure-linked downstream prediction.

ProteinSage remains competitive on fold classification and secondary-structure prediction, although modest gaps remain relative to the strongest baseline on these two tasks. These benchmarks emphasize complementary aspects of protein organization, including coarse-grained topology and short-range backbone patterns. Thus, the downstream results are consistent with the central design of ProteinSage: converting predicted residue-proximity relationships into pretraining objectives can improve transfer under reduced model and training budgets, without requiring structural inputs at inference.

### 2.4 ProteinSage scales efficiently under structural constraints

We then examined how ProteinSage scales with model capacity, data size and optimization budget under the same frozen-representation structural reasoning setting. Using a unified evaluation head and metric, this analysis separates the effects of parameter count, training data and training tokens from downstream fine-tuning variation.

As shown in Fig. 4a, increasing model size from 77 M to 650 M parameters yields consistent gains across all four benchmarks. Performance improves from 39.9 to 66.9 on CASP14, 43.2 to 65.4 on CASP15, 53.7 to 77.3 on CAMEO, and 0.385 to 0.556 on Recent. The 150 M model already recovers a substantial fraction of the total gain relative to the 77 M model, suggesting that structure-constrained pretraining uses additional capacity efficiently rather than relying only on overparameterization.

**Fig. 4.**
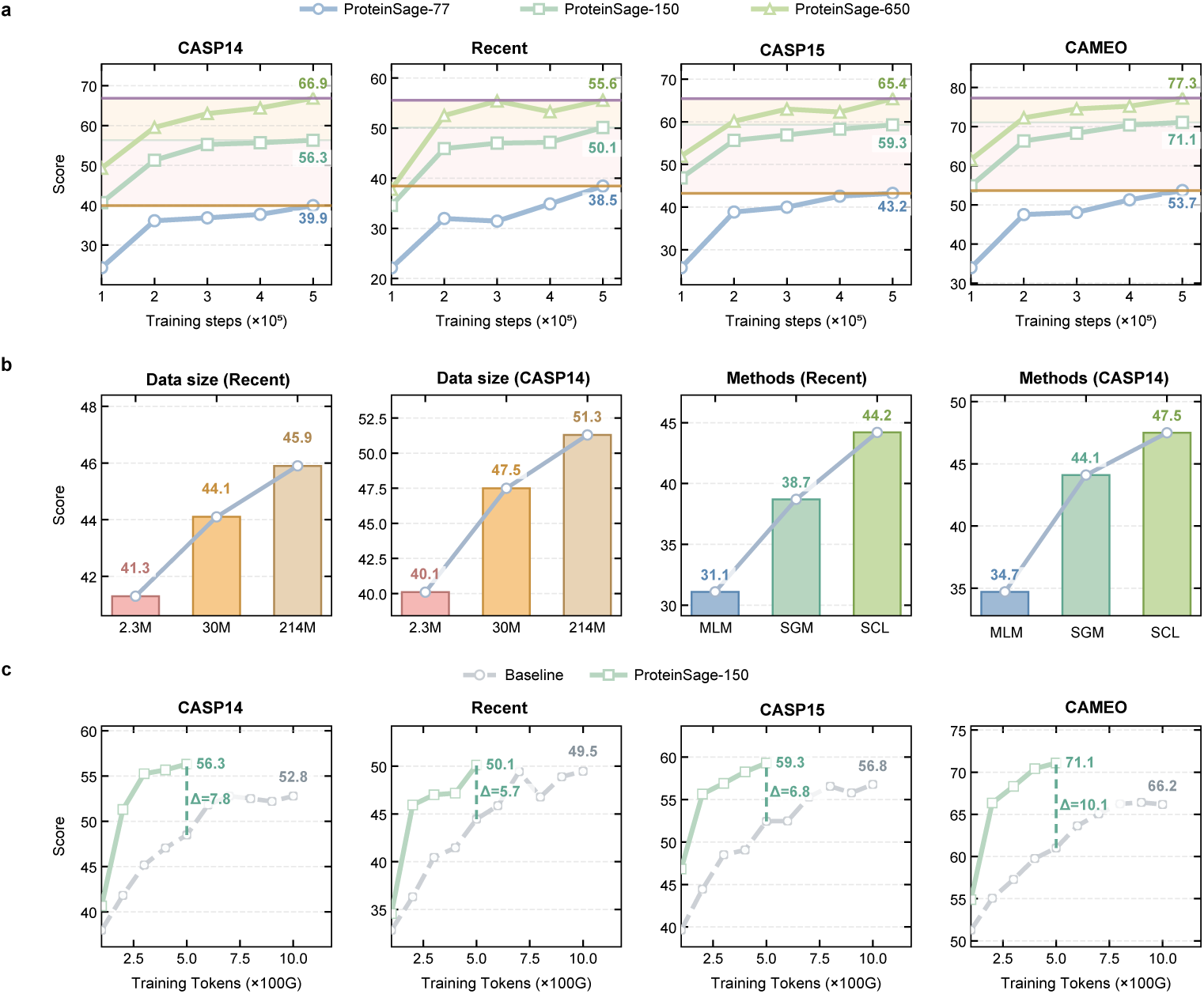
Scaling behavior and ablation analysis of ProteinSage on unsupervised contact-map prediction. **(a)**, Parameter scaling from 77M to 650M parameters yields monotonic performance gains, indicating efficient utilization of model capacity. All models are trained on the SageDB-Large dataset (214M sequences). **Left of (b)**, Data scaling analysis showing consistent performance improvements with increasing training data. **Right of (b)**, Learning-objective ablation demonstrating that the structural causal learning (SCL) objective outperforms the standard masked-language-modeling (MLM) objective and alternative variants. **(c)**, Performance as a function of training tokens (×100 G) compared with a conventional MLM baseline, showing faster convergence and a higher performance plateau for ProteinSage.

Data scaling shows a similar trend (Fig. 4b). Expanding the training set from 2.3 M to 214 M sequences improves CASP14 performance from 40.1 to 51.3 and Recent performance from 41.3 to 45.9, with smooth intermediate gains at 30 M sequences. These sub-linear but consistent improvements indicate that ProteinSage benefits from additional data while maintaining a favorable scaling regime under structural constraints.

Optimization curves further show rapid convergence (Fig. 4c). Across benchmarks, performance improves quickly during early training and begins to plateau after approximately 300 G training tokens. This indicates that the structure-constrained objective accelerates acquisition of contact-relevant features and allows training to be capped earlier without substantially sacrificing final accuracy. Together, these scaling results show that ProteinSage improves with both model capacity and data volume, while retaining high training efficiency under reduced token budgets.

### 2.5 Explicit structural objectives drive ProteinSage performance

Performance and efficiency gains in ProteinSage reflect how predicted structural priors are encoded into the pretraining objective, rather than the mere use of structural information. To test this, we compared three objectives under matched architecture, data and evaluation settings: (i) **MLM**, the masked-language-modeling objective implemented with standard random masking (SRM); (ii) **SGM**, structure-guided masking that preferentially selects spatially proximal but sequence-distant residue pairs while retaining token-level reconstruction; and (iii) **SCL**, structural causal learning that further converts these paired residues into directional dependency-prediction targets. All ablations use a 150 M-parameter ProteinSage model trained on SageDB-Large. For external reference, we follow the PSL [9] architecture and training recipe and train on the same corpus (approximately 900 M sequences, ∼4× our data).

The ablation shows a clear progression from MLM to SGM to SCL. On Recent (Fig. 4b), performance increases from 31.1 to 38.7 and 44.2, while on CASP14 it rises from 34.7 to 44.1 and 47.5. The improvement from MLM to SGM indicates that AFDB-predicted residue-proximity priors help identify more informative training positions, whereas the further gain from SGM to SCL shows that explicitly modeling the dependencies between these positions provides additional benefit. Thus, ProteinSage gains are driven by transforming structural relationships into learning targets, not simply by exposing the model to predicted structural information.

Training dynamics support the same conclusion (Fig. 4c). At 300 G tokens, SCL outperforms MLM by +10.1 on CASP14, +8.4 on CASP15, +11.0 on CAMEO and +6.6 on Recent. With 500 G tokens, SCL remains ahead of an MLM model trained on 1.0 T tokens by +3.5, +2.5, +4.9 and +0.6 points, respectively. Together, these results show that objective-level structural supervision accelerates learning and raises the performance ceiling under reduced token budgets.

Algorithmically, MLM must infer long-range dependencies indirectly from random token reconstruction. SGM introduces predicted residue-proximity priors by biasing which positions are masked, but its target remains token-level recovery. SCL further changes the optimization problem by requiring the model to predict directional dependencies between paired residues. This MLM–SGM–SCL progression demonstrates that ProteinSage improves pretraining efficiency by elevating predicted structural relationships from sampling cues to explicit objectives.

### 2.6 Evaluation of ProteinSage–Miner on metagenomic rhodopsin candidate identification

To evaluate whether ProteinSage representations support practical protein discovery, we applied ProteinSage–Miner to microbial type-I rhodopsins, a structurally conserved but sequence-diverse family of retinal-binding membrane proteins [42]. They are sequence-diverse retinal-binding membrane proteins with conserved seven-transmembrane topology, extracellular N-termini and TM7 retinal-binding lysines [43]. Their light-driven ion-transport activity and established optogenetic and bioelectronic utility [44–46] make them a useful benchmark for assessing whether a model can assign informative priorities to low-similarity candidates by integrating sequence, structure, and functional constraints.

We applied **ProteinSage–Miner** to metagenomic sequence screening and compared it with CNN, LSTM, Transformer and attention-based neural models following [47], as well as alignment-based approaches including BLAST [48] and MMseqs2 [49], and profile HMM-based domain annotation using PF01036 (Bac rhodopsin)[50]. Performance was evaluated in two settings: (i) a curated microbial rhodopsin benchmark assessing classification on held-out sequences; and (ii) large-scale screening of the Global Microbial Gene Catalog (GMGC) [51], where prioritized candidates were assessed using structural and domain-level criteria.

On the curated benchmark, ProteinSage achieves higher classification performance than the neural baselines (Fig. 5a–d), reaching an accuracy of 0.996 and an F1-score of 0.988, compared with 0.991 and 0.977 for ESM-2. These results indicate improved prioritization of rhodopsin-like sequences under matched supervised evaluation, rather than establishing discovery beyond all sequence-based methods.

**Fig. 5.**
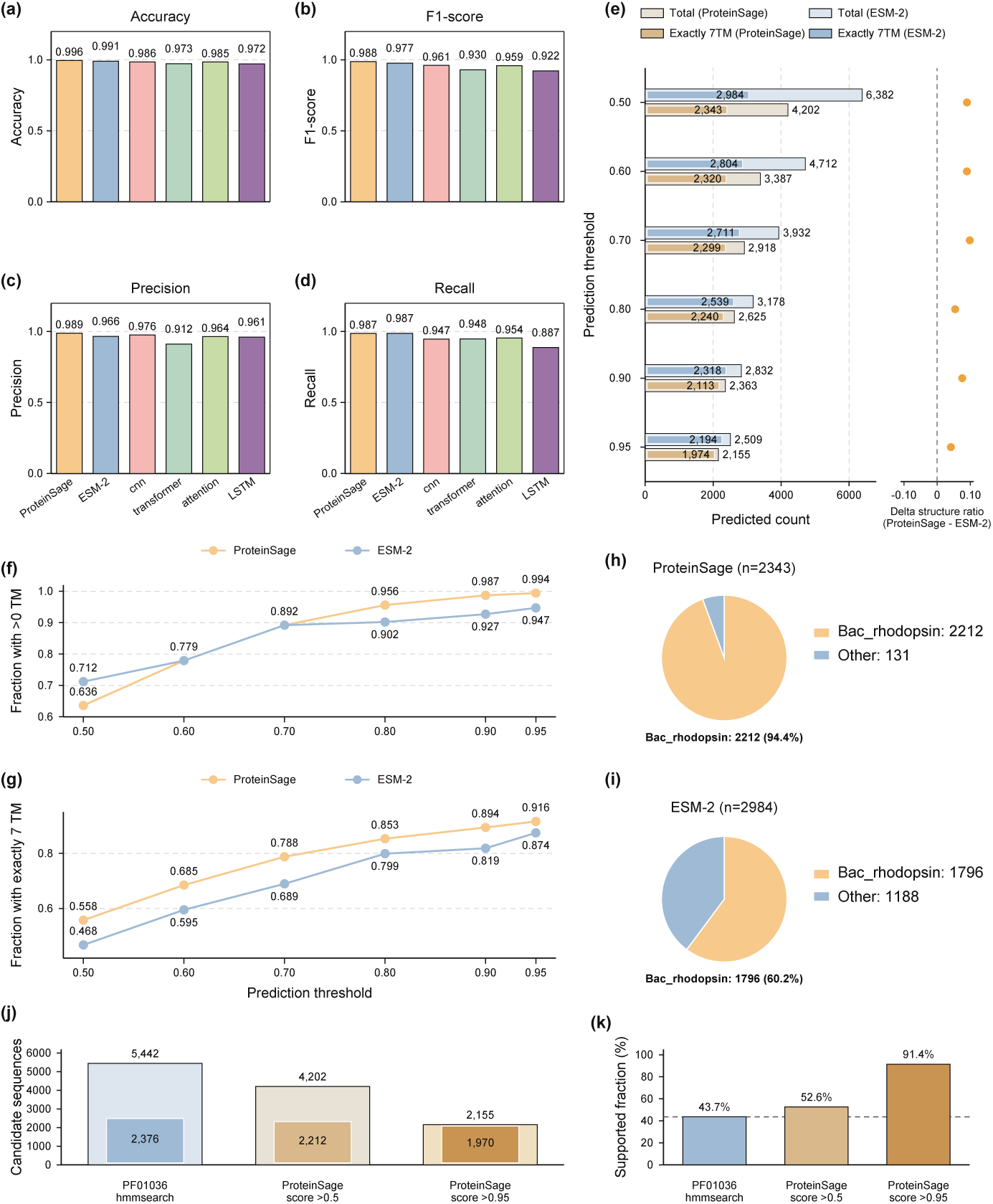
Comprehensive comparison between machine learning models, sequence-based models, and ProteinSage on protein mining tasks. **(a)–(d)**, Benchmark metrics for ProteinSage and representative baselines. **(e)**, Total candidates and candidates with exactly seven predicted transmembrane helices (7TM) at each confidence threshold; dots show the ProteinSage–ESM-2 difference in exact-7TM fractions. **(f,g)**, Fractions of candidates with at least one predicted transmembrane helix (f) or exactly seven (g), across confidence thresholds. **(h),(i)**, Domain composition of predicted sequences, showing an increased proportion of bona fide Bac_rhodopsin domains among ProteinSage predictions compared with ESM-2. **(j),(k)**, Relationship between ProteinSage ranking and PF01036 profile-HMM screening. ProteinSage-Miner substantially overlaps with PF01036 HMMER screening while additionally providing a model-based confidence score for candidate prioritization. Increasing the ProteinSage score threshold enriches candidates with both rhodopsin-domain support and seven-transmembrane topology, indicating the proportion microbial rhodopsin with high confidence significantly increase as the score threshold rises.

In GMGC screening, ProteinSage-ranked candidates show stronger enrichment for rhodopsin-associated structural features. Across confidence thresholds, a larger fraction of ProteinSage candidates satisfy seven-transmembrane constraints, with 55.8%–91.6% forming complete seven-transmembrane architectures, compared with 46.8%–87.4% for ESM-2 (Fig. 5e,g). Domain annotation further shows that ProteinSage predictions are enriched for Bac rhodopsin-like profile annotations: 2,212 of 2,343 predicted sequences (94.4%) carry a *Bac rhodopsin* annotations, compared with 1,796 of 2,984 (60.2%) for ESM-2, which shows a higher fraction of non-*Bac rhodopsin*or unassigned domains. Although domain hits can only serve as one piece of evidence for the candidate positive status of rhodopsin, this consistency check indicates that ProteinSage has a more accurate grasp of the sequence/structure features of rhodopsin compared to ESM-2.

Overlap analysis with sequence-based pipelines provides additional insight. ProteinSage recovers the majority of candidates identified by BLAST and MMseqs2, while identifying 534 candidates not retrieved by either pairwise-alignment method (Extended Data Fig. 4a). By contrast, ESM-2 identifies fewer candidates outside BLAST/MMseqs2 coverage (Extended Data Fig. 4b). Direct comparison shows that ProteinSage covers most ESM-2 predictions while contributing additional candidates (Extended Data Fig. 4c), suggesting broader candidate coverage while maintaining agreement with established sequence-based signals.

To compare ProteinSage ranking with profile HMM-based screening, we searched GMGC using the PF01036 HMM profile with HMMER (*E <* 1 × 10^−5^). This identified 5,442 hits, of which 2,376 (43.7%) were predicted by DeepTMHMM to contain seven transmembrane helices (Supplementary Data 3). At a score threshold of *>* 0.5, ProteinSage retained 4,202 candidates, including 2,212 sequences supported by both PF01036 annotation and seven-transmembrane topology, covering 91.8% (2,182 consecutive sequences) of the HMM-identified PF01036-positive seven-transmembrane set (Supplementary Data 2). As the ProteinSage score threshold increased, the total candidate number decreased substantially, whereas PF01036-positive seven-transmembrane candidates were largely retained; even at *>* 0.95, 1,970 of 2,155 candidates (91.4%) remained supported by both criteria (Fig. 5j,k). These results indicate that ProteinSage preserves broad agreement with profile HMM and topology-based screening while providing a global confidence score for prioritizing low-similarity candidates for downstream experimental validation.

### 2.7 Wet-lab validation identifies six previously uncharacterized microbial rhodopsins

Application of ProteinSage–Miner enabled the prioritization of candidate microbial rhodopsins from large-scale metagenomic data. Among fifteen model-prioritized candidates selected for experimental testing, six low-similarity sequences were experimentally confirmed as functional type-I proton pumps.

From 247 candidate type-I rhodopsin sequences identified by ProteinSage–Miner screening (Fig. 6a,b; Methods, Section 4.2.4), fifteen representatives were selected for validation (Supplementary Data 2; Extended Data Fig. 5). Heterologous expression in *Escherichia coli* showed that six candidates produced clearly coloured pellets, unlike the pale beige empty-vector control (Fig. 6c). Pigmentation, a hallmark of retinal-binding type-I rhodopsin expression, ranged from magenta or orange-red (q_927, q_1008, q_1831) to orange-yellow (q_706, q_467, q_1451), whereas the remaining candidates showed little or no detectable colour. These six sequences showed low global sequence identity to the curated rhodopsin reference set (38.9–48.3%), and their best-aligned UniProtKB sequences showed only 22.8–41.9% global sequence similarity (Supplementary Data 2), consistent with their phylogenetic distance from characterized family members.

**Fig. 6.**
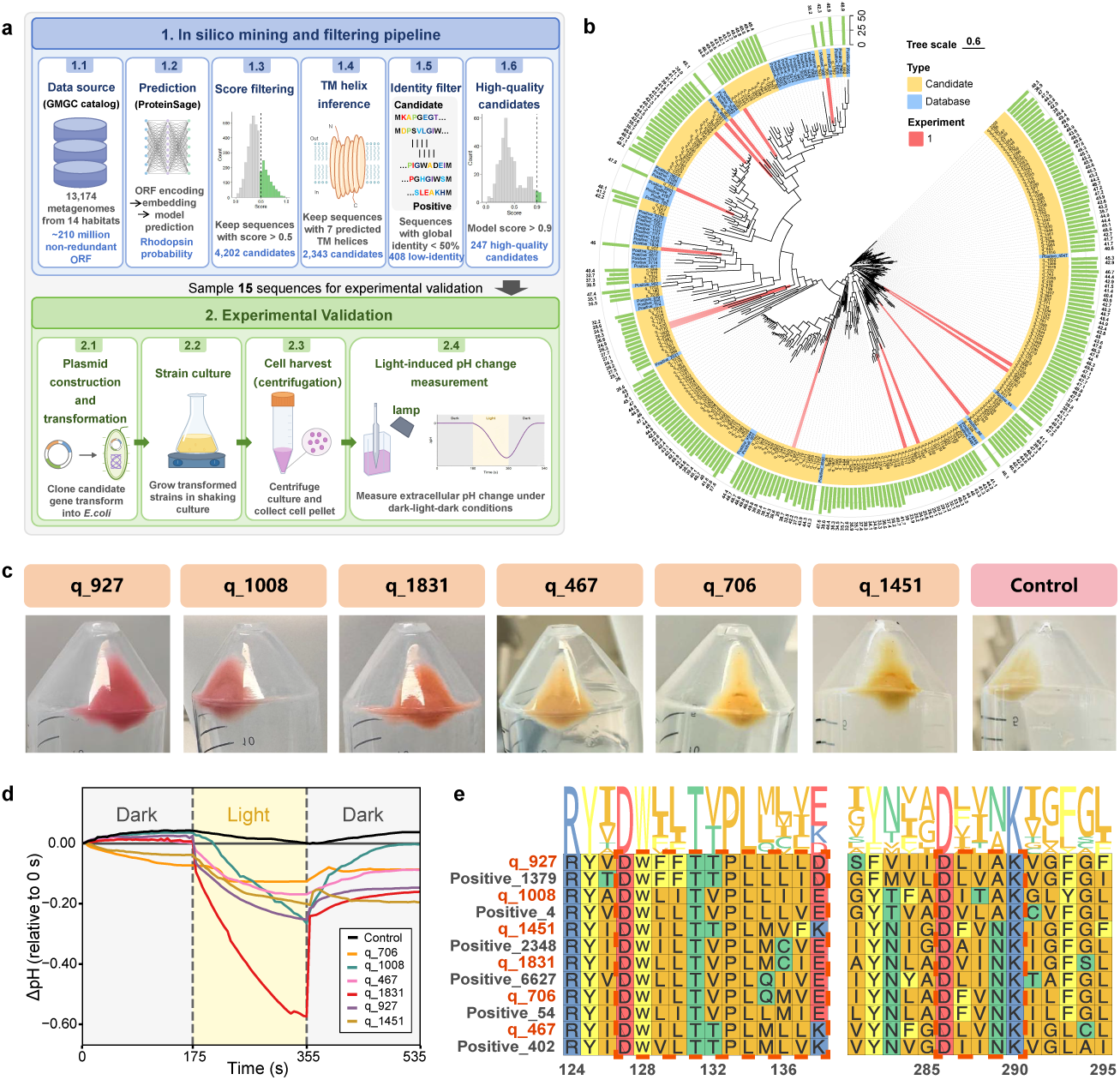
Wet-lab validation identifies six previously uncharacterized microbial rhodopsins. **(a)** Schematic overview of the microbial rhodopsin mining task. The GMGC was constructed from approximately 2.3 billion raw ORFs; the 90%-amino-acid non-redundant subcatalog used for screening (GMGC10.90nr.faa) contains 210,478,083 protein sequences. Candidate filtering and experimental validation are described in the Methods. **(b)** Maximum-likelihood phylogeny of 247 candidate sequences identified by ProteinSage together with their closest positive reference sequences (n = 56). Rings indicate wet-lab validation (red), sequence identity and source (sky blue, reference; yellow, candidate), and maximum pairwise sequence identity to the nearest reference. **(c)** Cell pellets of Escherichia coli expressing six rhodopsin candidate sequences selected from the 15 experimentally tested constructs, together with the empty-vector control. These six samples displayed distinct magenta-to-orange pellets compared with the control, indicating successful rhodopsin expression. Results for the remaining constructs are provided in Supplementary Data 5. **(d)** Light-dependent changes in external solution pH for the six coloured cell suspensions shown in (c), measured to assess proton-pumping activity. **(e)** Multiple sequence alignment of the six validated rhodopsins with their closest database homologs, showing conserved TM3 (“DxxxTxxxxxxD/E/K”) and TM7 (“DxxxK”) motifs.

Functional assays confirmed light-driven proton-pumping activity for all six coloured samples. Upon illumination, each induced acidification of the external unbuffered medium, with q 1831 showing the strongest response, whereas the empty-vector control showed no pH change (Fig. 6d; Supplementary Data 4). The presence of conserved motifs in TM3 (*DxxxTxxxxxxD/E/K*)[52] and TM7 (*DxxxK*)[53], which correspond to functionally constrained residues involved in proton transfer and retinal binding, respectively, further supported these results (Fig. 6e). Together, these results support the annotation of these sequences as previously uncharacterized type-I proton-pumping rhodopsins.

Overall, ProteinSage–Miner efficiently prioritized six functional microbial rhodopsins with low global sequence identity to known rhodopsins. Although the validation rate was moderate, the result is notable given the low sequence similarity of the candidates and the difficulty of prioritizing remote homologs from large metagenomic sequence space using conventional sequence-based screening alone[54]. Because heterologous expression of membrane proteins in *E. coli* is inherently variable, additional model-prioritized candidates may require further optimization to reveal activity[55]. These findings indicate that predicted-structure-informed protein language modeling provides useful sequence-only prioritization signals for experimental screening of low-similarity candidates, complementing conventional sequence- and profile-based searches in metagenomic protein discovery.

## 3 Discussion

ProteinSage establishes objective-level structure-constrained pretraining as a practical strategy for protein language modeling. While existing PLMs learn from large-scale sequence objectives [1–5] and recent structure-aware models incorporate structural modules, tokens, alphabets or multimodal representations [21, 23–25], ProteinSage uses structural information at a different level: AFDB residue-proximity priors are encoded into the pretraining objective rather than supplied as structural inputs at inference. Conceptually, this can be viewed as a restricted form of teacher-source structural prior transfer, in which AFDB/AlphaFold-derived predictions provide weak residue-level supervision for pretraining. This allows sequence-only inference while directing learning toward long-range residue dependencies during pretraining.

The key contribution is therefore not the use of structural information alone, but its conversion into explicit learning targets. The MLM–SGM–SCL ablation shows this progression: SGM improves over conventional MLM by selecting more informative residue pairs, whereas SCL further improves performance by directly modeling dependencies between those pairs. Together with the scaling analyses, these results indicate that objective-level structural supervision improves learning efficiency and representation transfer under reduced data and token budgets.

The use of AFDB also defines the scope of ProteinSage. AFDB greatly expands structural coverage [17, 18], but AlphaFold-derived structures may carry PDB coverage bias, MSA and homology dependence, single-conformation bias and geometric idealization [56, 57]. ProteinSage does not reconstruct atomic coordinates, complete distance maps or conformational states, nor does it replace experimental structural biology.

Thus, although it is conceptually related to knowledge distillation or teacher–student learning, it does not distill AlphaFold as a full structure-prediction model; instead, it transfers coarse residue-proximity constraints into a sequence model (Extended Data Fig. 1b). It is therefore less suited to problems dominated by extreme orphan folds, rapid evolutionary divergence, allosteric transitions or conformational dynamics, where static AFDB-derived priors may provide incomplete supervision.

Within this scope, ProteinSage provides a data-efficient route to transferable protein representations. Its gains across structural reasoning and supervised downstream benchmarks show that predicted residue-proximity priors can improve representation learning without increasing model size or requiring structural inputs during inference. The rhodopsin application further illustrates its use as a structure-aware, sequence-only prioritization strategy for metagenomic candidates, complementary to BLAST, MMseqs2, HMMER and Pfam rather than a replacement for sequence- or profile-based search.

More broadly, ProteinSage suggests that future protein foundation models need not rely solely on brute-force sequence scaling. By combining sequence-only inference with explicitly bounded structural prior transfer, this framework provides a basis for more data-efficient PLMs. Incorporating experimental structures, conformational ensembles, dynamics-aware priors and experimentally validated feedback may further extend this direction while preserving the practical advantages of sequence-only modeling.

## 4 Methods

### 4.1 ProteinSage

#### 4.1.1 Data curation

We constructed three different-sized pretraining sets, SageDB-Small, SageDB-Middle and SageDB-Large, based on the Foldseek-released AFDB clustering [58] (Extended Data Fig. 6a). We used Foldseek’s AFDB-50 clustering, which combines MMseqs2 sequence clustering at 50% identity and 90% overlap [49] with subsequent Foldseek structural clustering.

SageDB-Small contains one representative from each non-singleton structural cluster (≈ 2.27 M sequences). SageDB-Middle contains ≈ 30 M sequences obtained by resampling additional members around these cluster centroids to increase diversity while controlling redundancy. SageDB-Large corresponds to the full AFDB collection (≈ 214 M sequences). This three-tier design provides a consistent data-scaling ladder, from low redundancy to broader coverage, while keeping clustering criteria fixed across datasets. It allows us to isolate the effect of data volume from model capacity and optimization settings, and is used for the data-scaling experiments in Fig. 4b.

For the ESM-C comparison, we report the publicly reported counts of approximately 83 M UniRef, 372 M MGnify and 2 B JGI sequence clusters obtained at 70% sequence identity, totaling approximately 2.455 B clustered training entries and rounded to 2.5 B in the resource plots [31, 59].

#### 4.1.2 Structure-guided masking (SGM)

To enhance the model’s ability to learn structurally meaningful dependencies, we introduce the structure-guided masking (SGM) method.

We construct a residue–residue proximity graph from AFDB structures and use it to guide masking toward residues that are spatially close but sequence-distant. This targeted masking encourages the model to internalize non-local interactions that are essential for protein structure and function (Extended Data Fig. 6b). The spatial distance threshold of 6 Å is a conservative choice within the commonly used 4.5–8 Å range for contact-map definitions; prior large-scale evaluations report best fold-level discrimination for C*α*/C*β* cutoffs in the 6.5–7.5 Å band, while community practice often employs C*β* ≤8 Å for benchmarking [60, 61]—our stricter 6 Å setting aligns with this literature and is further validated by the ablations.

As shown in Extended Data Fig. 7a, this yields a fused mask that emphasizes non-local structural constraints through SGM while retaining broad token-level coverage through the 12% random MLM component. The spatial-proximity threshold and the SGM–MLM mixture ratio were not chosen heuristically, but were determined by ablation analysis (Extended Data Fig. 8a,b). Specifically, the 6 Å cutoff gave the best overall contact-prediction performance among 5 Å, 6 Å and 7 Å thresholds, balancing overly restrictive contact selection at 5 Å against noisier pair selection at 7 Å. Under a fixed 15% total masking rate, the 3% SGM + 12% MLM setting provided the strongest and most stable downstream performance compared with higher SGM ratios. We therefore used the 3%:12% split to introduce compact structure-guided pair-prediction targets while preserving sufficient base-sequence token reconstruction, thereby avoiding excessive sequence expansion and maintaining stable memory usage and throughput.

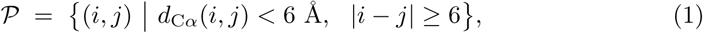

The overall strategy is as follows: let *K* denote the set of *key residues* that participate in any pair in *P*, i.e., *K* = {*i* | ∃ *j* : (*i, j*) ∈ *P*}. At each training step we:

1. recompute (and cache) *P* from the current structure to maintain independence of samples across steps;
2. uniformly sample without replacement a set of *pair anchors S* ⊆ *P* such that the set of distinct endpoint positions, *U*_key_ = U_(_*_s,t_*_)∈*S*_{*s, t*}, covers approximately 3% of the base-sequence positions. Both endpoints of each selected pair are included in the SGM mask set, with shared endpoints counted only once toward the masking budget; 3. additionally sample 12% *random*residues from the complement K to preserve general token-level difficulty and keep the total mask rate at 15% (matching our BERT-like baseline [62]).

All masking rates are defined relative to the base-sequence length *n*. The disjoint SGM and random MLM mask sets cover approximately 3% and 12% of these positions, respectively, giving a total base-sequence masking rate of 15%.

Operationally, the procedure can be viewed as sampling from a sparse proximity graph *G* = (*V, E*) where *V* indexes residues and *E* = *P*. We exclude short-range pairs (|*i* − *j*| *<* 6) to avoid over-representing local secondary-structure regularities already captured by standard MLM. By concentrating masking on *K* while maintaining a modest random component, the model is repeatedly exposed to structure-informed residue dependencies without sacrificing coverage of generic sequence statistics, which we find accelerates convergence and improves transfer on structure-linked endpoints.

#### 4.1.3 Structural causal learning (SCL)

To further enhance the model’s ability to capture structure-aware dependencies, we build on SGM and introduce the structural causal learning (SCL) objective. Rather than modifying how SGM identifies key residue pairs, SCL operates on top of it: given a known source residue, the model is encouraged—through a causal prediction pathway—to infer its spatially proximate neighbors. Concretely, SCL appends a compact pair-prediction trailer to each sequence and enforces directed attention from the source residue(s) to the appended target token(s), enabling explicit learning of residue-to-neighbor relationships.

On a length-*n* sequence we first apply base masking with **MLM** and **SGM**. From the SGM-derived structure pairs 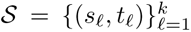 we then build a *pair-prediction trailer* by appending **two** slots per pair: a *source* slot 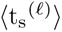 and a *target* slot 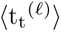. The augmented sequence and length are described in Eq. 2.

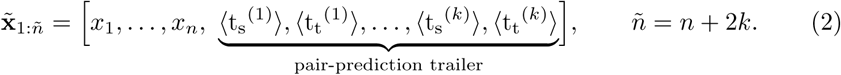

Each trailer pair encodes a *directed* dependency: 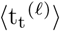 must be predicted *causally* from its partner 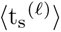 and the base context. As shown in Eq.3, to avoid directional and positional bias, we randomly swap the roles within each pair at every step.

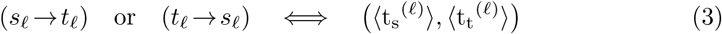

As shown in Extended Data Fig. 7a, we enforce a simple masking scheme that keeps the base sequence bidirectional, lets each trailer token attend to the base, and permits trailer-to-trailer attention *only* within the same pair and only in the directed (t_s_ →t_t_) direction:

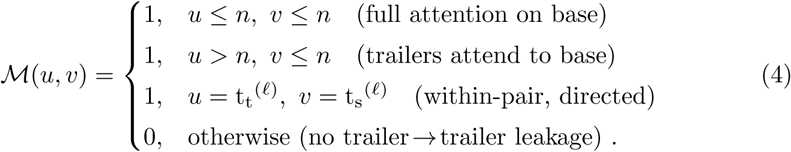

Thus, information flows from the selected source residue (via its trailer ⟨t_s_⟩ and the base) to the corresponding target trailer ⟨t_t_⟩, while trailers from different pairs remain isolated. Here, *k* = |*S*| denotes the number of selected residue pairs. The 3% SGM masking rate refers to |*U*_key_|*/n*, rather than to *k/n*. Each selected pair contributes one source slot and one target slot to the trailer. Thus, the augmented sequence length is *n* + 2*k*, and the relative increase in sequence length is 2*k/n*. The appended trailer slots are not included in the base-sequence masking-rate calculation.

In Extended Data Fig. 6b, beyond point-to-point (1 → 1), SCL can form small *span* groups where a set of sources predicts a short contiguous target span (*n* → *m* with 1 *< n, m <* 6). Implementation follows the same trailer construction, appending *m* target slots per group and allowing directed attention from the group’s source slot(s) to the target span (left-to-right within the span). In practice, the main results use the 1 → 1 setting for the best accuracy–efficiency trade-off.

Base-sequence masking remains unchanged: SGM masks approximately 3% of the original sequence positions, and random MLM masks an additional, disjoint 12%, giving a total masking rate of 15%. SCL appends source and target slots for the selected pairs without changing these base-sequence masking rates. Pair directions are re-sampled at each step to prevent trivial memorization. Ablations show that the 1 → 1 variant offers the best accuracy–efficiency balance, while span SCL broadens relational context at a modest cost in sequence length. In sum, SCL converts a small subset of structure-critical sites into explicit, directed dependencies while keeping computational overhead bounded.

#### 4.1.4 Architecture

To keep comparisons transparent, we adopt a widely used Transformer encoder backbone with engineering changes and expose our contributions entirely through the masking policy and learning objective. All ProteinSage variants use pre-normalization with *LayerNorm*, rotary positional embeddings (RoPE), multi-head self-attention, and a GELU-activated feed-forward network. Unless otherwise noted, dropout is disabled, sequences are truncated/padded to 1,024 tokens, and training runs in bfloat16 with ZeRO optimization and activation checkpointing.

We instantiate three capacities to probe scaling: *77M*, *150M*, and *650M* parameters. The small model (77M) uses 28 layers with *d*_model_=480, 20 attention heads, and a feed-forward dimension of 1280. The middle model (150M) uses 30 layers with *d*_model_=640, 20 attention heads, and a feed-forward dimension of 1707. The large model (650M) uses 33 layers with *d*_model_=1280, 20 attention heads, and a feed-forward dimension of 3424. The SGM and SCL objectives are layered on this backbone.

In Fig. 4c, the baseline follows the PSL[9] 150 M setting: a masked language modeling (MLM) objective trained on ≈ 900 M sequences with a total budget of 1 trillion tokens. Our reimplementation reproduces the final performance reported in PSL, confirming the fidelity of our data, training schedule, and evaluation pipeline, and validating the comparisons presented here.

#### 4.1.5 Optimization and training schedule

The optimization and training schedules differ among the three models. We tailor the learning-rate schedule by model size.

For the **ProteinSage-77M** and **ProteinSage-650M** models we use Adam (*β*_1_=0.9, *β*_2_=0.95, *ɛ*=10^−8^) with a cosine decay from a peak learning rate of 2×10^−4^ to a floor of 4×10^−5^. For the **ProteinSage-150M** model we use the same optimizer with a cosine decay from 5.5×10^−4^ to 5.5×10^−5^. All runs employ a token-based warmup of 2.5% and gradient clipping at 1.0, with a global batch size of 1,024. To enable fair comparison, the total pretraining budget is fixed across sizes at 5.0×10^11^ tokens (500 G tokens). Training is distributed with DeepSpeed (ZeRO stage 1) and activation partitioning; tensor and pipeline parallelism are set to 1 for all experiments.

#### 4.1.6 Training objective

We adopt a unified “recover the masked token” formulation that accommodates all three components—**MLM** (implemented with standard random masking, **SRM**), **SGM**, and **SCL**—within a single encoder and softmax head. On a length-*n* input, let K denote key residues. At each step we form two disjoint mask sets on the *base* sequence:

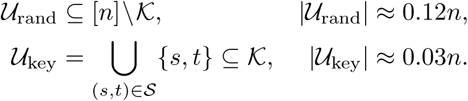

so the total base mask rate remains 15%. The SGM loss is averaged over distinct masked base-sequence positions, so a position shared by multiple selected pairs contributes only once to this loss. These yield two token-reconstruction terms that differ only in *where* the masked positions are drawn:

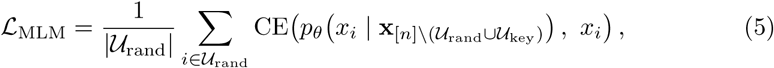

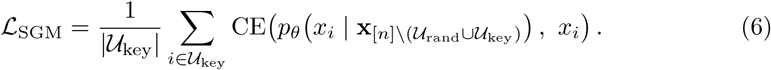

SCL operates on an *appended* trailer tied to key pairs. For the sampled directed pairs 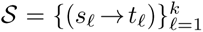 (point-to-point) or grouped spans (*n* → *m*), let *T* be the set of appended trailer targets (size *k* for 1→1, or Σ*_g_m_g_* for spans). We supervise these trailer tokens causally, conditioning on the base context and the corresponding source residue(s):

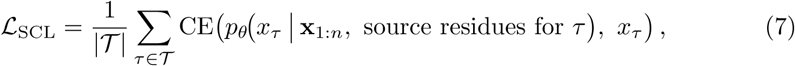

with the attention mask permitting trailer→base and (within-pair) source→target flow while blocking trailer↔trailer leakage.

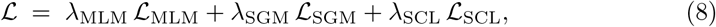

Total objective. The full loss is a weighted sum with *λ*_MLM_=*λ*_SGM_=*λ*_SCL_=1 unless noted otherwise. This formulation keeps implementation simple (a single decoder head), isolates the effect of the masking *policy* (MLM vs. SGM) from the causal *targeting* (SCL), and allows clean ablations and scaling under matched compute.

### 4.2 ProteinSage-Miner

#### 4.2.1 Dataset construction

To construct a positive dataset of microbial rhodopsins, we compiled 183 experimentally validated type-I microbial rhodopsin sequences from UniProt[63] and the published literature. These seed sequences were used to expand the dataset via BLASTp v 2.15.0[64] (E-value cutoff 1e-5) searches against UniProt, Pfam[50], NCBI, KEGG[65], EggNOG[66] and MicRhoDE[67]. Predicted transmembrane helices were inferred with DeepTMHMM[68], and sequences were filtered to remove duplicates and any entries lacking seven predicted transmembrane helices, yielding a final positive set of 4,068 sequences (Supplementary Data 1). Pfam[50] domain annotation indicated that all positive sequences contained the PF01036 (Bacteriorhodopsin-like domain).

For the negative dataset (see Data Availability section), we retrieved proteins from UniProt that share similar physicochemical properties, cellular localization, or apparent functional roles with rhodopsins but are not type-I microbial rhodopsins. These included non-microbial photoreceptors, G protein-coupled receptors (GPCRs), membrane proteins with comparable numbers of transmembrane helices, transporters, and a diverse set of functionally unrelated enzymes (e.g., hydrolases and oxidoreductases). The diversity of the negative set was chosen to improve classifier robustness and generalizability. Because the average length of the rhodopsin positive sequences was 261 amino acids, only negative sequences of 160–360 aa were retained. These negative sequences were clustered with CD-HIT v4.8.1[69]. To avoid performance inflation from homologous sequences, we performed a cluster-based split: the dataset was first clustered at 70% sequence identity, and the resulting clusters—rather than individual sequences—were divided into training, validation, and test sets in an 8:1:1 ratio.

#### 4.2.2 Architecture and training setting

As shown in Extended Data Fig. 7b, we fine-tuned the ProteinSage (650M) backbone using an adapter-based strategy. ProteinSage-Miner employs a six-layer **Transformer adapter** followed by a classification head. During training, only the adapter and classifier parameters were updated, while the backbone parameters of **ProteinSage**—used as the base model—were kept frozen. This adapter-based design allows efficient specialization for rhodopsin recognition while preserving the pre-trained biological priors of the foundation model.

Optimization was performed using the AdamW optimizer with a learning rate of 2 × 10^−5^ and a cosine learning rate scheduler. Training was run for one epoch with a batch size of 64 and a maximum sequence length of 1,024. Data loading was parallelized with 4 workers, and validation was conducted once per epoch.

#### 4.2.3 Training objective

For rhodopsin classification, the adapter maps each input sequence representation to a binary prediction score. The model is trained with a binary cross-entropy loss to distinguish rhodopsins from non-rhodopsin membrane proteins:

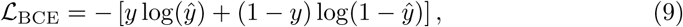

where *y*^ is the predicted rhodopsin probability and *y* ∈ {0, 1} is the ground-truth label. Only the lightweight adapter and classification head are optimized, while the pretrained ProteinSage backbone remains frozen.

#### 4.2.4 Mining pipeline

ProteinSage-Miner was applied to the GMGC to mine putative type-I microbial rhodopsin sequences. The GMGC was constructed from approximately 2.3 billion raw ORFs derived from 13,174 metagenomes representing 14 distinct habitats. For large-scale inference, we used the publicly available 90%-amino-acid non-redundant subcatalog GMGC10.90nr.faa, which contains 210,478,083 protein sequences. All sequences in this screening set were submitted to ProteinSage for feature extraction; the resulting embedding vectors were processed by an adapter module and a classification head to yield binary predictions for rhodopsin candidates. Candidate sequences were ranked by model prediction confidence score, and those with score *>* 0.5 were kept (*n* = 4, 202). Predicted transmembrane helices were then inferred using DeepTMHMM[68], and only sequences with seven predicted transmembrane helices were retained (*n* = 2, 343).

To obtain a conservative candidate set for downstream characterization, we further applied sequence-based filtering against a curated positive reference set of known type-I rhodopsins. Candidate sequences were first compared with the positive reference set using BLASTp[64] and MMseqs2 [49] to identify close sequence matches and nearest reference proteins. Pairwise global sequence alignments were then performed using EMBOSS Needle[70] to calculate final sequence identity values. Candidates sharing ≥ 50% sequence identity with any positive reference sequence were considered high-similarity candidates and excluded, whereas candidates with *<* 50% identity to all positive references were retained. This filtering resulted in a final set of 408 low-sequence-similarity candidate type-I rhodopsins. Among them, 247 sequences with model scores greater than 0.9 are regarded as high-quality candidates (Fig. 6a).

A phylogenetic tree was constructed for the 247 candidates together with their most similar positive reference sequences (*n* = 56). Multiple sequence alignment was performed with MAFFT v7.490[71] (–auto), alignments were trimmed with trimAl v1.4.1[72] (-automated1) to remove spurious sequences and poorly aligned regions, and a maximum-likelihood phylogeny was inferred using FastTree 2 v2.1.11[73] (Fig. 6b). Fifteen representative sequences were selected for experimental validation. These sequences had an average sequence identity of 37.9% to the positive reference set and an average best pairwise identity of 50.79% among candidates.

As an additional sequence-based baseline, we evaluated profile HMM–based sequence profiling using HMMER v3.3.2[74] against the Pfam database[50]. The Bac rhodopsin family PF01036 was used as the HMM-based rhodopsin reference family. In contrast to pairwise alignment approaches such as BLAST and MMseqs2, profile HMMs capture family-level conservation patterns through probabilistic sequence profiles, providing a complementary benchmark for detecting remote rhodopsin homologs.

#### 4.2.5 Experimental validation

All reagents were of analytical grade or higher and were used as received unless otherwise stated. 15 candidate type-I microbial rhodopsin sequences were synthesized (Wuhan GeneCreate Biological Engineering Co., Ltd.) and cloned into the pET21a(+) expression vector using the NcoI and XhoI restriction sites. Chemically competent *Escherichia coli* C41(DE3) cells were used as the expression host. A total of 15 constructs carrying candidate sequences and one empty-vector control were transformed and cultured in 20 mL lysogeny broth (LB) supplemented with ampicillin (100 mg L^−1^) at 37^◦^C with shaking.

When the culture reached an OD_600_ of ≈0.6, protein expression was induced by addition of isopropyl *β*-D-1-thiogalactopyranoside (IPTG; final concentration 1 mM) and all-trans retinal (final concentration 10 *µ*M) under control of the T7 promoter with lac operator (T7-lacO). Cultures were incubated overnight at 37^◦^C with shaking at 120 rpm, and cells were harvested by centrifugation (2,520 × *g*, 15 min, 20^◦^C). Cell pellets were washed three times with physiological saline (0.9% NaCl), and pellet coloration was recorded by visual inspection (Fig. 6c).

Expression of rhodopsin typically produces a visibly colored pellet (orange to magenta to blue)[43], with color intensity roughly correlating with expression level; pellets from non-expressing or weakly expressing cultures appear pale yellow to tan, as observed for the empty-vector control. Six constructs that produced significant or a certain amount of magenta/orange pellets (q 927, q 1008, q 1831, q 706, q 467 and q 1451) were selected for functional testing of proton-pumping activity.

For functional characterization, proton transport activity was measured by monitoring light-dependent changes in the pH of an unbuffered NaCl solution, following previously established protocols[75, 76]. Briefly, cell pellets were resuspended in 5 mL of unbuffered 100 mM NaCl. A pH electrode (AQUASEARCHER™ AB41PH, OHAUS, USA) was inserted into the suspension, and measurements were performed in a dark environment. Illumination was provided by a 300 W xenon lamp controlled by a shutter system.

After a 3-min dark equilibration period to stabilize the solution pH, samples were illuminated for 3 min, followed by a 3-min dark recovery period. pH was recorded at 5-s intervals throughout the experiment. A light-induced decrease in the external pH (acidification) that reverses during the subsequent dark period was interpreted as evidence of outward proton transport by the expressed rhodopsin.

#### 4.2.6 Visualization

Phylogenetic trees were visualized using TVBot (https://chiplot.online/tvbot.html)[77], and multiple sequence alignments using ggmsa[78]. Structures were predicted with AlphaFold3 [79] and visualized using the AlphaFold Server viewer (Extended Data Fig. 5) or ChimeraX 1.8[80] (other figures).

## 5 Data Availability

The downstream datasets, labels and structural files supporting this study are publicly available in Zenodo [81]. The structural pretraining data were derived from protein clusters available through the Foldseek cluster resource at https://cluster.foldseek.com/. The PF01036 Hidden Markov Model (HMM) profile used for rhodopsin sequence profiling can be obtained from the InterPro/Pfam entry at https://www.ebi.ac.uk/interpro/api/entry/pfam/PF01036?annotation=hmm. The publicly available metagenomic protein database used for large-scale model inference, GMGC10.90nr.faa (210,478,083 protein sequences), can be downloaded from https://gmgc.embl.de/download.cgi.

## 6 Code Availability

The ProteinSage source code and example usage instructions are publicly available at https://github.com/biomap-research/ProteinSage under tag v1.0.1 and archived as version v1.0.1 in Zenodo [82]. The pretrained model checkpoint weights are publicly available at https://huggingface.co/biomap-research/ProteinSage.

## 7 Acknowledgements

We thank BioMap for providing access to its high-performance computing infrastructure and its engineering team for technical support and infrastructure optimization for large-scale concurrent model training. Bioinformatic support from the High-performance Computing Platform of Peking University is also gratefully acknowledged.

## 9 Author Contributions Statement

J.N., X.Z. and T.L. (Tianhong Li) conceived and supervised the study, acquired funding, provided resources, and contributed to the interpretation of the results and revision of the manuscript. L.S. and L.C. contributed to the study design, algorithm development, data analysis and interpretation, and preparation and revision of the manuscript. T.L. (Tang Liu) and Q.L. contributed to data collection and curation, as well as bioinformatic and computational analyses. G.Z. and H.W. contributed to engineering optimization for large-scale model training, benchmark construction and model evaluation. T.L. (Tang Liu) and X.D. performed the protein-based wet-lab experimental validation. All authors discussed the results, reviewed the manuscript and approved the final version.

## Extended Data

**Extended Data Fig. 1.**
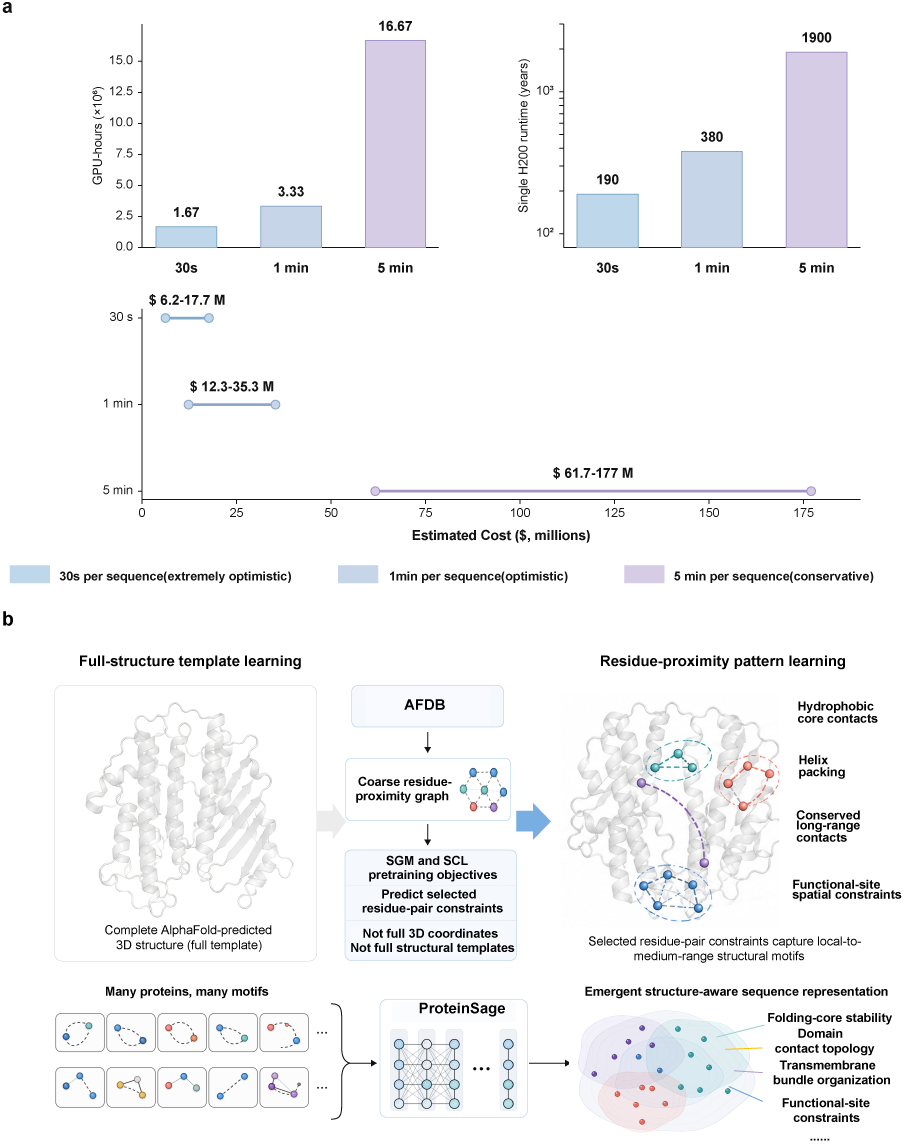
Computational motivation and scope of AFDB-derived structural priors in ProteinSage. **a**, Estimated computational burden of structure-dependent metagenomic-scale screening. ProteinSage uses predicted structures only during pretraining to construct learning signals, whereas downstream tasks and large-scale screening require only amino acid sequences as input. In the non-redundant GMGC screening set used in this study, approximately 200 million protein sequences were evaluated. Even under an optimistic estimate of one minute of GPU time per medium-length protein for AlphaFold2 structure prediction, predicting structures for all candidates would require approximately 200 million GPU minutes, or 3.3 million GPU hours, equivalent to about 380 years of continuous computation on a single GPU. This estimate excludes additional costs from MSA search, CPU preprocessing, I/O, failed runs and structural file storage. **b**, ProteinSage learns local residue-proximity regularities rather than complete structural templates. AFDB-predicted structures are used only to derive selected residue-pair proximity constraints, such as spatially adjacent but sequence-distant contacts within local domains, packing regions or functional-site neighborhoods. ProteinSage does not use full three-dimensional coordinates as fold templates and is not trained to memorize complete predicted structures. Instead, repeated exposure to partial residue-dependency motifs allows the model to learn transferable sequence representations informed by local and long-range structural constraints.

**Extended Data Fig. 2.**
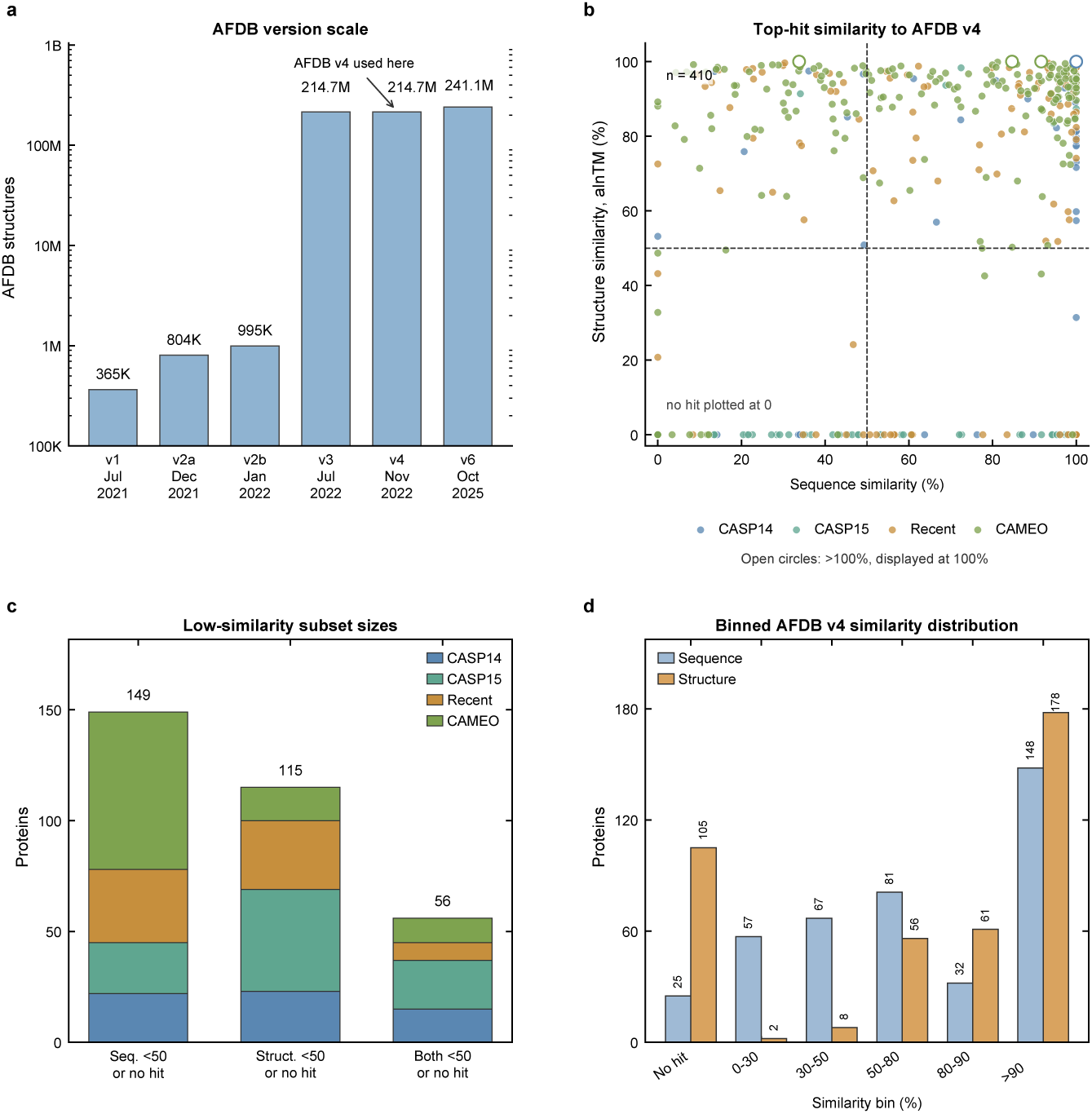
AFDB v4 similarity analysis of the structural reasoning datasets (CAMEO, CASP15, CASP14 and Recent). **a**, Evolution of AlphaFold Database scale across major releases, highlighting AFDB v4 as the database version used for this comparison. **b**, Per-protein sequence and structure similarity to the top AFDB v4 hit. Sequence similarity is measured by strict global identity, defined as exact global-alignment matches divided by the maximum of query and target lengths. Structure similarity is measured by Foldseek strict top-hit alnTM. Four reported alnTM values slightly exceeded 1 (1.002–1.003); these are displayed at 100% as open circles for visualization only. Uncapped values are retained in the Source Data and used for threshold-based classification; this display adjustment does not change subset membership or binned counts. Dashed lines indicate the 50% thresholds; proteins without AFDB hits are plotted at zero. **c**, Sizes of conservative low-similarity subsets, where no-hit proteins are counted as below 50%. Among 410 proteins, 149 have sequence similarity below 50% or no sequence hit, 115 have structure similarity below 50% or no structure hit, and 56 satisfy both criteria. **d**, Binned distribution of sequence and structure similarity to AFDB v4, showing that most proteins have high structural similarity, while a non-trivial subset remains distant by sequence, structure, or both. Overall, the analysis identifies a conservative subset of 56 proteins with low similarity to AFDB v4 under both sequence and structural criteria, supporting leakage-controlled evaluation on AFDB-distant proteins.

**Extended Data Fig. 3.**
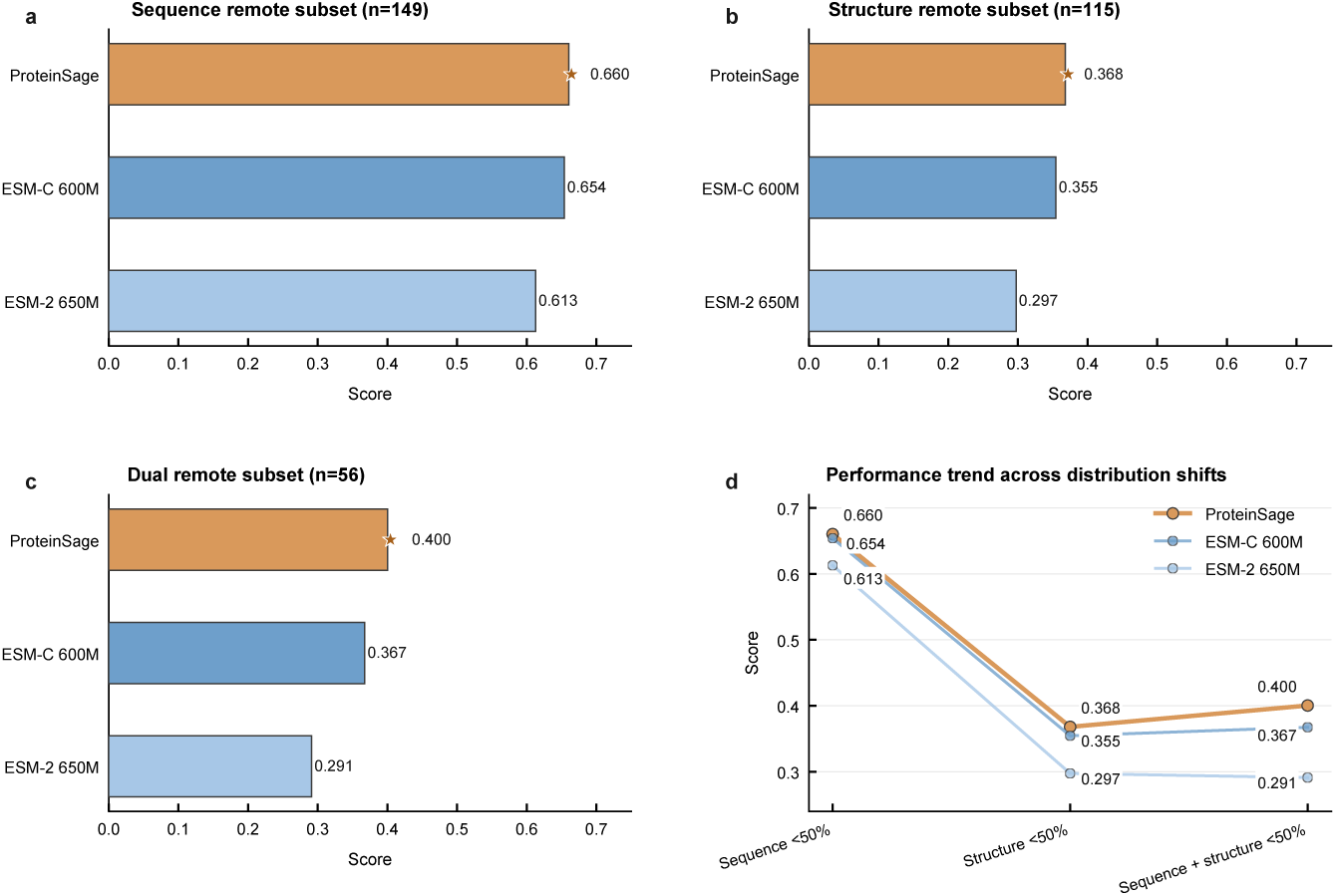
Contact-map prediction on low-similarity test subsets. **a,** Performance on proteins with low sequence similarity to the training set (*n* = 149). **b,** Performance on proteins with low structural similarity to the training set (*n* = 115). **c,** Performance on proteins with both low sequence similarity and low structural similarity (*n* = 56). In **a–c**, horizontal bars show the performance of ProteinSage, ESM-C 600M and ESM-2 650M, and stars indicate the best-performing model in each subset. **d,** Performance trajectories of the same models across the three subsets, showing how contact-map accuracy changes as the test distribution shifts from sequence-level novelty to structural novelty and combined sequence–structure novelty. ProteinSage achieves the highest Top L/5 precision across all three subsets.

**Extended Data Fig. 4.**
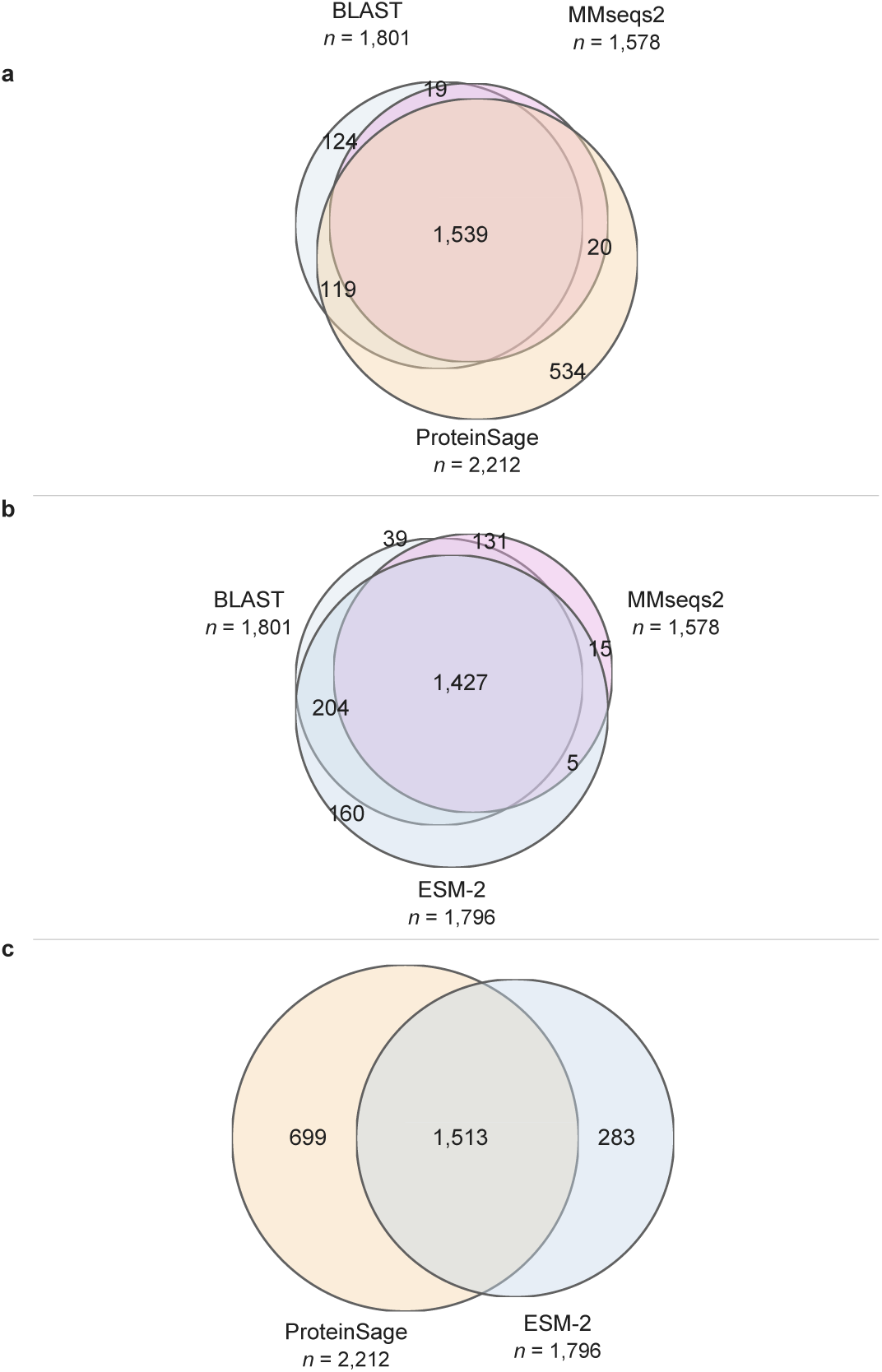
Overlap analysis of candidate sequences identified by ProteinSage, ESM-2 and alignment-based search methods. ProteinSage recovers most candidates identified by conventional sequence-search methods while also extending coverage to an additional set of method-specific candidate sequences. **a,** Venn comparison among BLAST, MMseqs2 and ProteinSage candidate sets, showing that the majority of candidates are shared across methods, whereas ProteinSage additionally contributes a distinct set of candidates not retrieved by the pairwise alignment pipelines. **b,** Venn comparison among BLAST, MMseqs2 and ESM-2, illustrating the overlap between conventional sequence-search approaches and the protein language model baseline. ESM-2 recovers a subset of candidates overlapping with BLAST and MMseqs2 but contributes fewer method-specific candidates. **c,** Direct comparison between ProteinSage and ESM-2 candidate sets. ProteinSage covers most of the ESM-2-supported candidate space while also identifying a larger ProteinSage-specific subset, suggesting improved candidate expansion and prioritization relative to the baseline protein language model. Together, these comparisons indicate that ProteinSage complements alignment-based screening by providing an additional structure-informed ranking signal for candidate prioritization.

**Extended Data Fig. 5.**
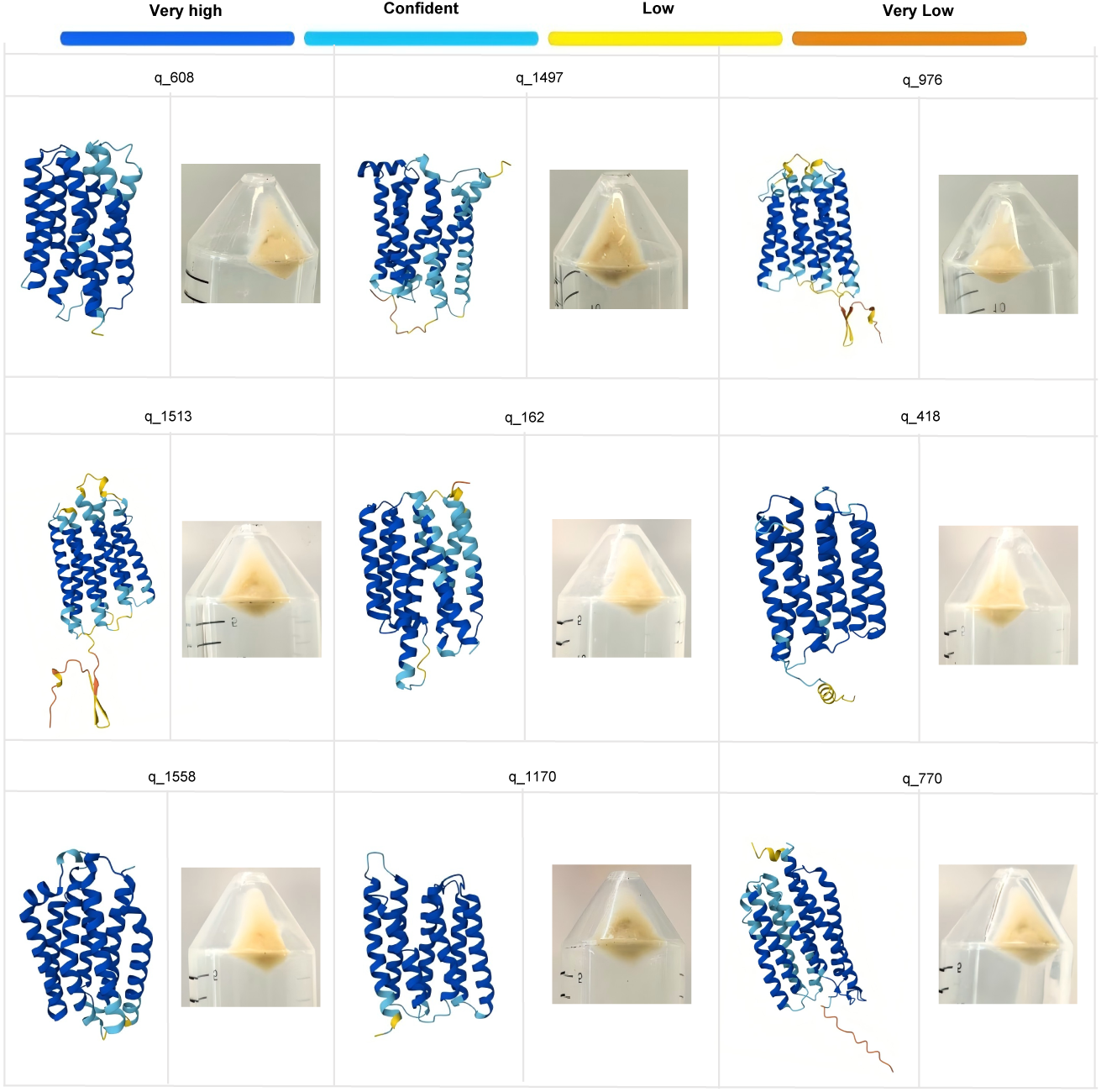
Predicted three-dimensional structures of nine sequences that were not successfully validated by wet-lab experiments. For each candidate, the AlphaFold 3-predicted structure is shown alongside the corresponding *Escherichia coli* cell pellet after heterologous expression. The models are coloured by local predicted local distance difference test (pLDDT) confidence: very high (dark blue, pLDDT *>* 90), confident (cyan, 70 *≤* pLDDT *≤* 90), low (yellow, 50 *≤* pLDDT *<* 70), and very low (orange, pLDDT *<* 50). All structures exhibit a seven–transmembrane-helix architecture characteristic of microbial type I rhodopsins. The transmembrane cores are predominantly assigned very high or confident pLDDT values, whereas lower-confidence regions occur mainly in terminal or loop segments. The corresponding pellets showed little or no detectable rhodopsin-associated pigmentation.

**Extended Data Fig. 6.**
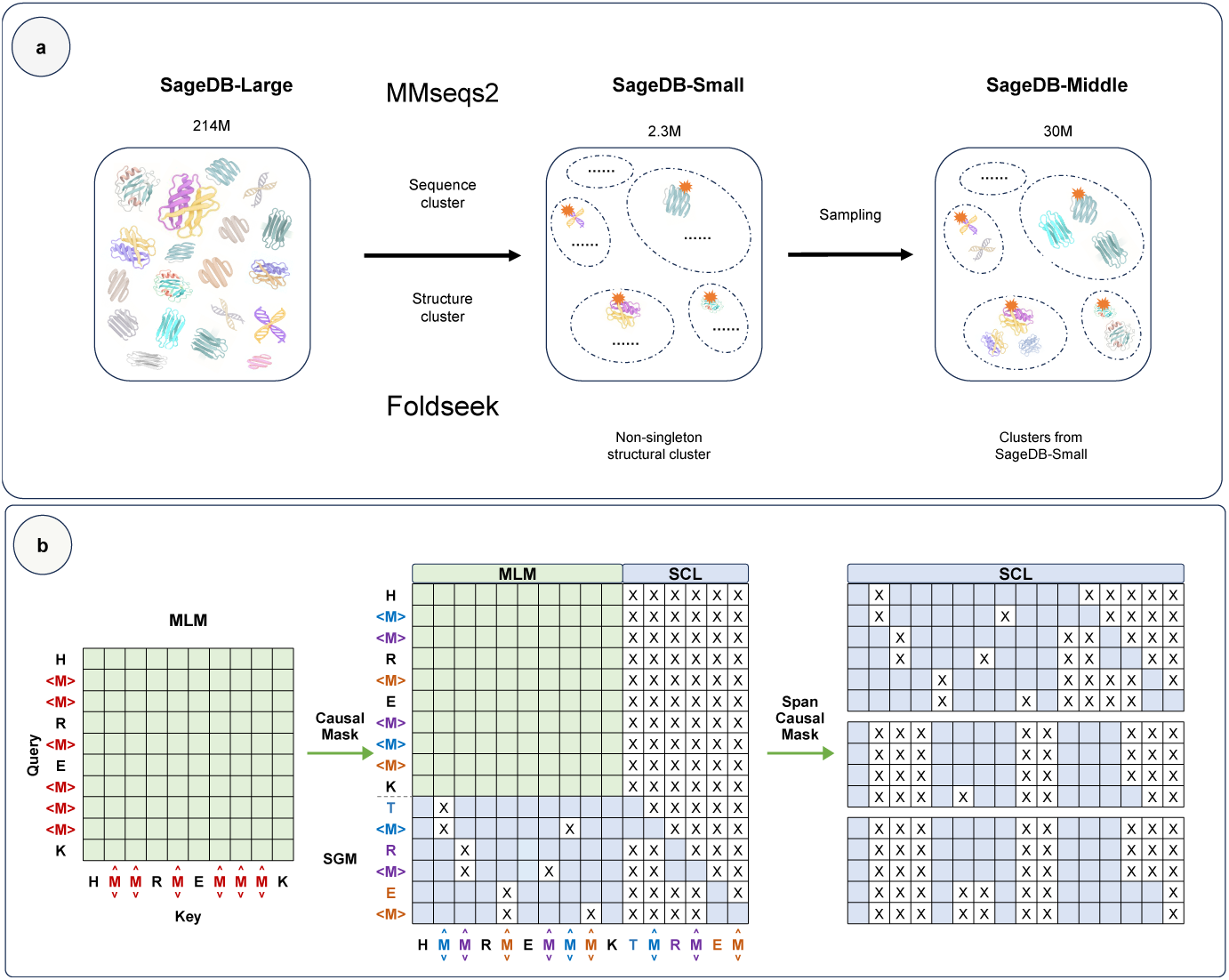
SageDB construction and objective-level structure-constrained pretraining in ProteinSage. **a**, SageDB curation and sampling pipeline. AFDB protein sequences are clustered using MMseqs2 and Foldseek. After fragment removal and filtering to non-singleton structural clusters, SageDB-Small samples one representative per cluster, SageDB-Middle resamples additional members around these centroids, and SageDB-Large uses the full AFDB collection. **b**, masking schemes and causal pair prediction. **Left**, the masked-language-modeling (MLM) objective uses standard random masking (SRM) and residue reconstruction. **Middle**, MLM with structure-guided masking (SGM) and point-to-point structural causal learning (SCL; 1 *→* 1). **Right**, span SCL with grouped *n → m* prediction (1 *< n, m <* 6).

**Extended Data Fig. 7.**
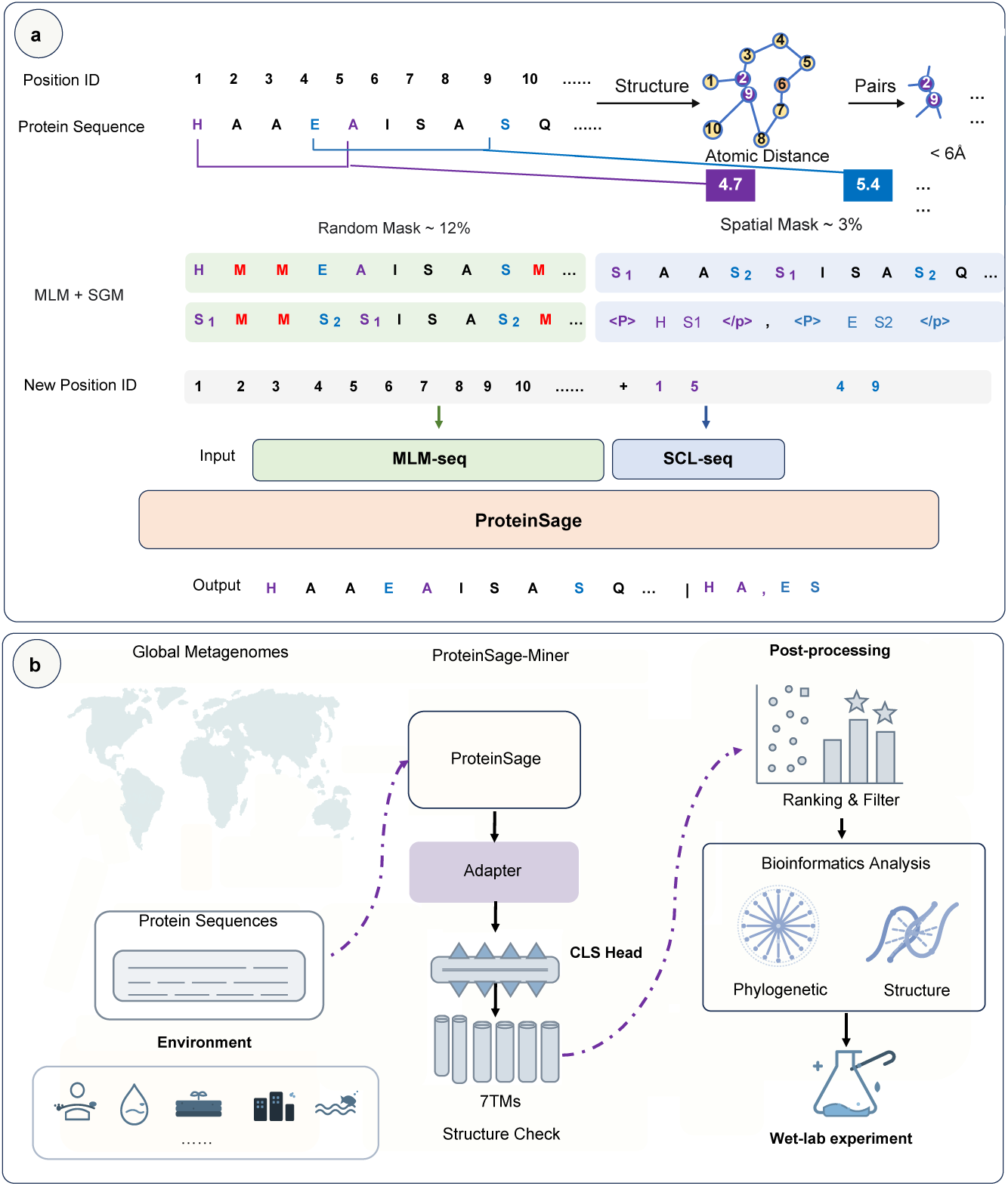
ProteinSage pretraining and ProteinSage-Miner discovery workflow. **a**, ProteinSage pretraining. Random and structure-guided masking generate augmented masked sequences, which are processed by the encoder and SCL trailer to jointly recover masked residues and predict paired trailer targets. **b**, ProteinSage-Miner for microbial rhodopsin discovery. Metagenomic proteins from 14 global environments are screened, filtered for seven-transmembrane topology, ranked by model confidence, and refined by domain and phylogenetic analyses before experimental validation.

**Extended Data Fig. 8.**
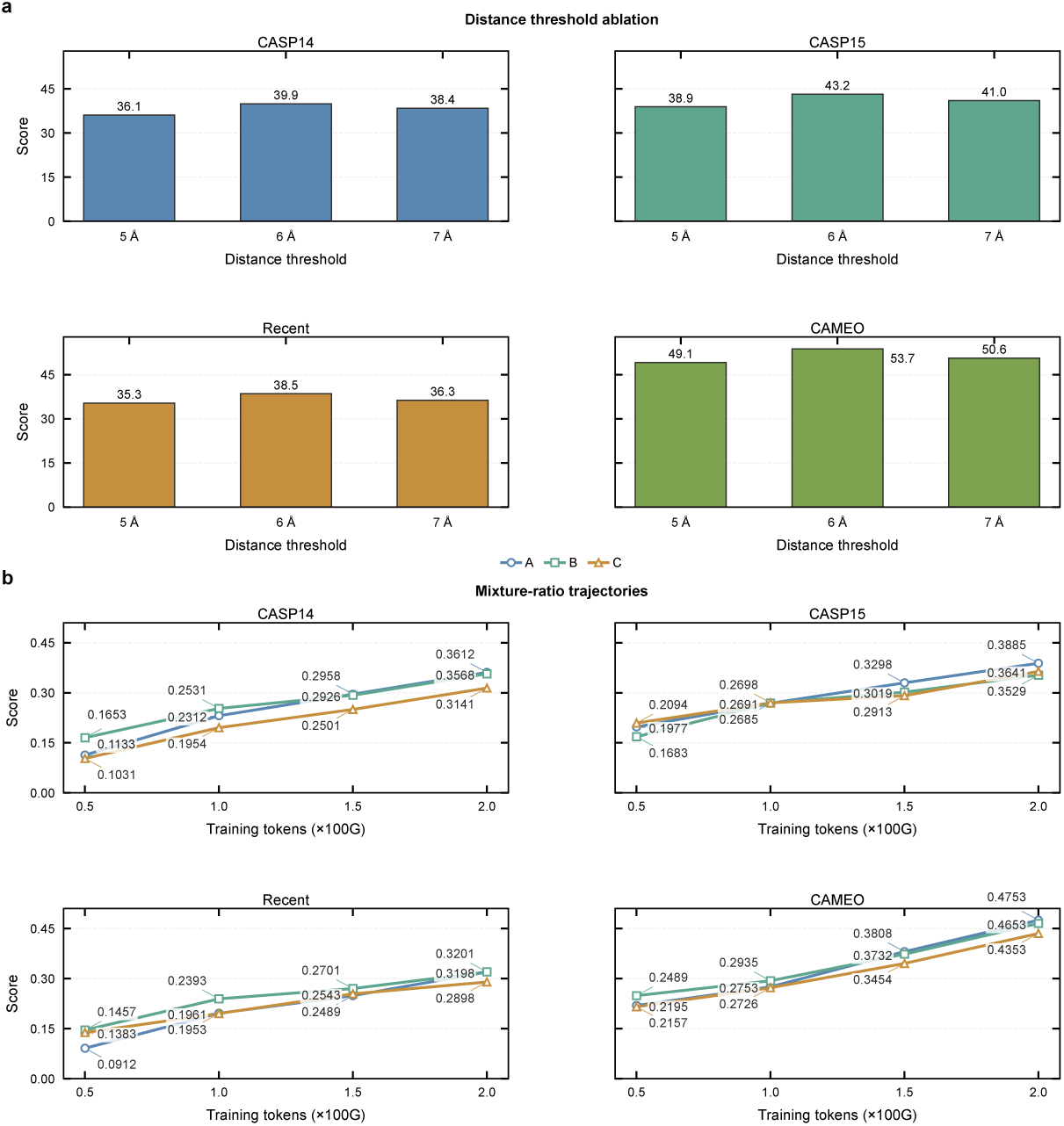
Ablation of ProteinSage structural-prior settings. **a**, Ablation of the spatial-proximity threshold used to select residue pairs for SGM/SCL training. ProteinSage-77M was trained on SageDB-Large and evaluated on CASP14, CASP15, Recent and CAMEO. Among 5 Å, 6 Å and 7 Å cutoffs, 6 Å gives the best overall performance, suggesting a balance between overly restrictive contact selection at 5 Å and noisier, less specific pair selection at 7 Å. **b**, Training-token trajectories for the SGM–MLM mixture-ratio ablation under a fixed 15% total masking rate. ProteinSage-77M was trained on SageDB-Large for 200 G tokens using three settings: A, 3% SGM + 12% MLM; B, 5% SGM + 10% MLM; and C, 7.5% SGM + 7.5% MLM. Moderate SGM ratios provide the strongest and most stable performance, whereas the highest SGM ratio degrades accuracy, likely because excessive pair-prediction targets shift optimization away from base-sequence token reconstruction. We therefore use the 3% SGM + 12% MLM setting as the default throughout this study.

